# Structural divergence in N-terminal domains of AAA proteases paraplegin (SPG7) and FtsH indicates a key structural function in complex formation

**DOI:** 10.64898/2026.04.22.720153

**Authors:** James G. Hyatt, Neil G. Paterson, Juliette M. Devos, Cristiano L.P. Oliveira, Christian M. Jessen, Sylvain Prevost, Andreas Hofmann, Jan Skov Pedersen, Anja Winter

## Abstract

AAA proteases are hexameric ATP-dependent metallopeptidases that perform crucial proteolytic activities within prokaryotic and eukaryotic membranes. Structurally, protomers are comprised of catalytically active C-terminal domains that are anchored to the membrane by an N-terminal autonomous folding unit. In this study, we determined the fold, stability, and oligomeric state of the N-terminal intermembrane domains of human spastic paraplegia type 7 (SPG7)/ paraplegin protein and its bacterial orthologue FtsH using circular dichroism (CD), small-angle X-ray scattering (SAXS), small-angle neutron scattering (SANS) and X-ray crystallography. Solution-state analysis revealed that the N-terminal domain of paraplegin is a monomer in solution whereas FtsH forms a dimer. Unexpectedly, the N-terminal domain of paraplegin presents as a domain-swapped homodimer in our crystal structure that involves the first helix and first two beta-strands from one monomer and beta-strand 3, helix 2 and beta-strand 4 from another symmetry-related molecule. However, together they form an assembly which is similar to protomers observed for the N-terminal regions of FtsH and AfG3L2. Drawing from our structural data, we postulate that domain-swapping interactions of the N-terminal regions contribute to stability of the AAA protease hexamer containing paraplegin, demonstrating the extensive flexibility of the N-terminal portion of this protein and its role in achieving the appropriate molecular architecture required for function.

**Graphical abstract:** 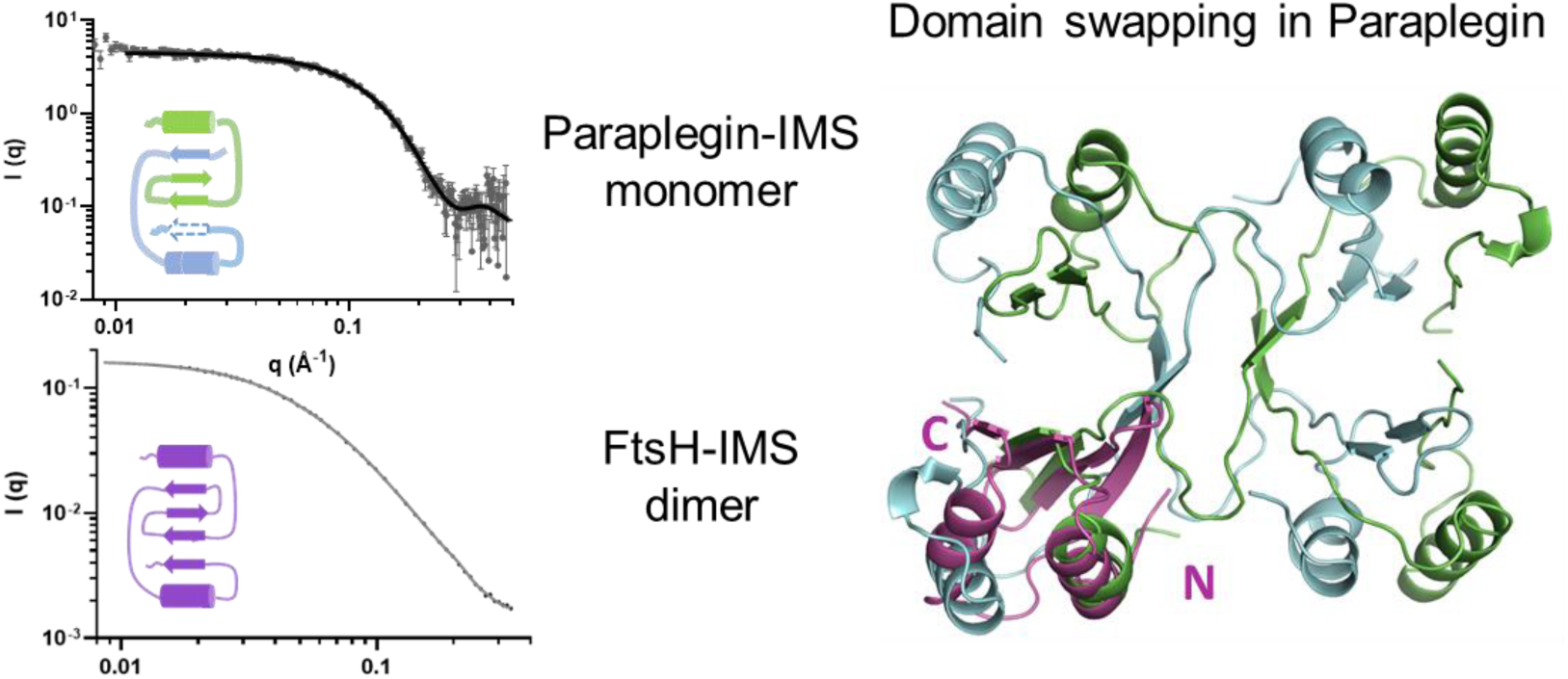

**Highlights:**

- FtsH-IMS forms a homo-dimer in solution, whereas paraplegin-IMS presents as a well-folded monomer in solution
- paraplegin-IMS crystallises as a domain-swapped homo-dimer but its domain-swapped monomers are structurally similar to other IMS-regions
- AfG3L2/paraplegin hexamer formation may be supported by domain swapping in paraplegin-IMS
- domain-swapping in paraplegin could be a Bonafide feature under certain cellular conditions and may be related to disease in spastic paraplegia

## 1 Introduction

The ATPases Associated with diverse cellular Activities (AAA+)-family of proteins are well conserved throughout animal and plant kingdoms, as well as in bacteria, carrying out crucial roles in cellular homeostasis. Dysfunction of AAA-proteases leads to wide-ranging and severe disruption to cellular function. In bacteria, deletion of the bacterial AAA-protease FtsH is lethal in *E. coli* [1] and causes severe growth defects in *B. subtilis* [2], supporting a crucial role for AAA protease in maintaining cellular homeostasis and viability. In eukaryotes, mitochondrial AAA proteases are responsible for mitochondrial proteostasis which is essential for regulation of the mitochondrial proteome and associated cellular functions, including subcellular protein trafficking, proteolysis of damaged, unfolded, or superfluous proteins, and the removal of damaged mitochondria [3–6]. Disruption of their functions has been implicated in pathologies such as cancer [7] and neurodegenerative disorders [8, 9]. Paraplegin, a human mitochondrial *m*AAA-protease, has originally been discovered and named for its involvement in hereditary spastic paraplegias (HSP) [10, 11], which most commonly manifest in progressive spasticity and weakness of the lower limbs.

At the molecular level, AAA-proteases are mostly homo-hexameric complexes, such as the FtsH proteases in bacteria [12, 13], however, some orthologues can differ in their subunit composition. FtsH-like AAA+ proteases in mitochondria are either matrix-facing (*m*-AAA proteases) or facing the inter-membrane space (*i*-AAA proteases), providing the possibility of functional interaction between assemblies [14]. In human mitochondria, *m*AAA protease hexamers may exist as homo-oligomeric complexes of the subunit Afg3L2, or hetero-oligomeric assemblies of both Afg3L2 with paraplegin/SPG7 [15]. In yeast, two *m*AAA-protease subunits, Yta10 and Yta12, which are orthologs to Afg3L2 and SPG7, respectively, have been identified [16] and their hetero-hexamer was shown in a 12 Å-resolution cryo-electron tomography (CET) map [17]. The variability amongst species in the oligomeric state and assembly of AAA proteases may suggest that the configuration of the assembly itself harbours subtle differences which contribute to substrate specificity and functionality, meriting closer examination [18]. For instance, a variable and tissue-specific subunit composition of mitochondrial *m*-AAA protease complexes was linked to hereditary spastic paraplegia [15] and thus directly associating complex composition with functionality.

Individual *m-*AAA protease subunits harbour two N-terminal transmembrane helices, which anchor the hexameric proteases to the inner mitochondrial membrane, as in the case for Afg3L2 and paraplegin [19], yeast Yta10 and Yta12 [16], and plant AtFtsH3 and AtFtsH10 [20], or to the periplasmic membrane as in the case for bacterial FtsH [21, 22]. The catalytic C-terminal ATPase and metallo-peptidase domains then protrude to the mitochondrial matrix or cell cytosol, respectively [22–25]. Structural information is available for the C-terminal ATPase and protease domains of FtsH from *E. coli* (PDB code 1LV7 [13]), *Thermatoga maritima* (PDB code 3KDS [26]), and human AfG3-L2 (PDB 6NYY [18]), all of them have been crystallised or resolved with cryo-EM in both their nucleotide-bound and apo states. Additionally, the three-dimensional structure of the functional cytosolic hexameric complex was later solved by Suno and colleagues in 2006 [27] with an outer diameter of 120 Å to 135 Å.

In contrast, the N-terminal region of AAA-proteases has attracted less attention, despite reports that deletion of the N-terminal region of FtsH and related AAA protease subunits impairs function [17] and reduces the formation of homo-oligomers [28]. This severely restricts degradation of membrane-embedded substrates [29], suggesting a key role for the N-terminal transmembrane domains in oligomer formation and protease function. Current studies into the topology of the N-terminal domain have shown two N-terminal transmembrane (TM) helices encasing a small ∼70-residue intermembrane space domain (IMS) which protrudes into the intermembrane space between the two mitochondrial membranes (inter-membrane space) or the periplasm [23, 30, 31]. Structural information for this domain can be obtained from crystal structures of FtsH from *Thermatoga maritima* (PDB code 4M8A) and *E. coli* (PDB code 4V0B [32]), FtsH NMR structure (PDB code 2MUY [32]), and AfG3-L2 NMR structure (PDB code 2LNA [33]), which show a mixed fold of two helices and four anti-parallel beta-strands. However, structural information for the N-terminal domain of the paraplegin orthologue is unavailable. Structural data of this region, and comparison with architectural features of paraplegin’s orthologues, would deepen our insight into the structure-function relationship and substrate recognition by AAA proteases; a critical requirement for understanding diseases such as hereditary spastic paraplegia.

Here, we set out to compare *in solution* states of the N-terminal domains of bacterial FtsH and human paraplegin to gain further insights into the structurally conserved N-terminal domains of the AAA-proteases FtsH and paraplegin. Both proteins were recombinantly produced and analysed using circular dichroism (CD), small-angle X-ray scattering (SAXS), small-angle neutron scattering (SANS), and X-ray crystallography. FtsH-IMS forms a homo-dimer in solution, supported by SAXS analysis, whereas paraplegin-IMS presents as a well-folded monomer in solution, but unexpectedly crystallises as a domain-swapped homo-dimer. Although the observed crystal structure may be influenced by high protein/salt concentrations and low pH during crystallization, it nevertheless offers a plausible model for AAA protease hexamer formation, where N-terminal domain-swapping dimerization facilitates oligomerization and may contribute to complex stability and substrate specificity.

## 2 Materials and Methods

### 2.1 Bioinformatic analysis and multiple-sequence alignment

Sequences from *m*AAA-protease subunits of FtsH (*E.coli*), Yta10 and Yta12 (yeast), AFL3G2 and paraplegin both from mouse and human were aligned using ClustalW [34]. Secondary structures were obtained either from crystal structures (FtsH ATPase domain, PDB accession number 1LV7, and peptidase domain, PDB accession number 2DI4) and/or PSIPRED [35] secondary structure predictions. The multiple sequence alignment was adjusted to preserve structural features and conserved residues in the ATPase domain and protease domain using Jalview [36]. Remaining N-terminal sequences were compared at both the primary and predicted secondary structure levels, and sequences manually aligned using Jalview [36] to obtain highest similarity.

### 2.2 Protein expression in *E. coli*

#### Paraplegin-IMS

A synthetic gene was generated encompassing the nucleotide sequences encoding for paraplegin intermembrane space region (amino acids 163 to 249, henceforth called Para-IMS) with an N-terminal 6x-histidine tag and TEV cleavage sequence into pRSET_A (Genewiz), and transformed into *Escherichia coli* BL21(DE3)pLysS cells (#70236-3, Novagen). Cells were grown in LB media supplemented with ampicillin (100 µg/ml, #BP1760-25, Fisher) and chloramphenicol (50 µg/ml, #B20841, Alfa Aesar), metal mix [37], and expression induced at an OD_600_ of 0.6 using 0.5 mM isopropyl-b-D-thiogalactopyranoside (IPTG) for 24 hours at 30 °C. Cells were harvested via centrifugation at 3500 g for 30 minutes, the pellet resuspended in PBS, and frozen at −20 °C.

#### FtsH-IMS

For cloning of the FtsH-IMS region into pRSET_C (amino acids 20-97, henceforth called FtsH-IMS), genomic *E.coli* DNA was isolated from DH5α cells (Invitrogen, #18265-017) and purified by combining two QIAGEN protocols (for MIDI prep and for isolation of genomic DNA). Briefly, the cell pellet was resuspended in 3.5 ml buffer B1 with 10 µl RNase A solution (100 mg/ml) by vortexing. A 100 µl aliquot of lysozyme stock solution (100 mg/ml) was added, and the mix was incubated at 37 °C for 30 minutes. After adding 1.2 ml of buffer B2, contents were mixed by inverting the tube several times and by vortexing for a few seconds and then incubated at 50 °C for 30 minutes. The QIAGEN MIDI prep column (Qiagen) was equilibrated with 10 ml of buffer QBT before the sample was applied. The column was washed two times with 10 ml of buffer QC, and the genomic DNA was eluted with 5 ml of buffer QF. Afterwards, the DNA was precipitated by adding 3.5 ml (0.7 volumes) room-temperature isopropanol. The tube was inverted 10 to 20 times, and the DNA spooled using a curved glass rod. The end of the rod containing the DNA was immediately transferred into a microcentrifuge tube, and 300 µl of a 10 mM Tris-HCl, solution pH 8.5, was added. After dissolving the DNA overnight on a shaker at room temperature, the microcentrifuge tube was centrifuged at 13000 g for 10 minutes at 4 °C and the dissolved DNA transferred into a clean microcentrifuge tube and stored at –20 °C.

Isolated DNA was used as a template for PCR to amplify DNA encoding *E. coli* FtsH (amino acids 20-97) with the following primers (MWG, Germany): FW 5’-CC GCT CGA GAT CAG AGC TTT GGG CCC AGC GAG-3’ (XhoI site) and RV: 3’-GGC GGA CTT CTT GGT TCG ATC CTT AAG GCC-5’ (EcoRI site). PCR was performed using *Taq* DNA polymerase (Promega UK Ltd.), purified via gel electrophoresis (1% agarose gel), and the appropriate DNA bands excised and extracted using the gel extraction kit from Qiagen following the manufacturers protocol. After restriction digest of amplicon and pRSET_C vector using XhoI and EcoRI (New England Biolabs as per manufacturer’s protocol) the products of digestion were purified via gel electrophoresis and gel extraction. Digested PCR product and plasmid were ligated using T4 DNA ligase (NEB), as per manufacturers protocol), and the ligation mix was transformed into XL1 Blue cells (Stratagene). Successful plasmid construction was verified via analytical digestion and sequencing (Big Dye Term Mix, DNA sequencing facility at the School of Biology, The University of Edinburgh, UK).

The IMS region of FtsH was expressed in BL21(DE3) cells (Invitrogen) in 7 L LB media (containing 100 µg/ml ampicillin) and induced when OD_600_ exceeded 1 with 0.5 mM IPTG (final concentration), overnight. Cells were harvested at 3000 g, 4 °C for 20 minutes, the cell pellet resuspended in buffer D1 (100 mM NaCl, 1.5 mM EDTA, 5 mM benzamidinium chloride, 1 mM PMSF, 0.1% Triton X-100, 20 mM Tris, pH 8.0) and stored at –20 °C.

### 2.3 Protein Expression in *E. coli* in matchout deuterated minimal medium

For the production of matchout-deuterated paraplegin-IMS protein, a kanamycin-resistant plasmid was constructed using the EZ-Tn5™ <KAN-2> Insertion Kit (EZ1982K, Lucigen®, Middleton, WI, USA). Protein production was carried out in the Deuteration Laboratory of the Institut Laue-Langevin (D-Lab, ILL, Grenoble, France) as described in Dunne et al., 2017 [38, 39]. *E. coli* BL21(DE3) competent cells (Invitrogen) were transformed with the kanamycin-resistant plasmid. Bacterial cultures were slowly adapted to growth in 85% deuterated minimal medium. A deuterium-adapted preculture was used to inoculate 1.2 L of matchout deuterated minimal medium in a 3 L bioreactor (Infors AG, Switzerland). A fed-batch fermentation culture was carried out at 30 °C until an OD_600_ of 17.5, protein expression was then induced with 1 mM IPTG. The culture was grown to a final OD_600_ of 19 after 21h of induction at 30°C. Cells were harvested by centrifugation and were frozen at -80 °C for long-term storage. The cell paste yield was 53 g wet weight from a 1.7 L culture.

### 2.4 Protein purification

#### Paraplegin-IMS

Para-IMS was purified from lysate after high-speed centrifugation to remove unbroken cells using Profinity IMAC resin (#156-0123, Bio-Rad) and eluted using an imidazole gradient consisting of buffer A (50 mM Na phosphate buffer pH 7.5, 200 mM NaCl, 10 mM imidazole) and buffer B ( 50 mM Na phosphate buffer pH 7.5, 200 mM NaCl, 250 mM imidazole). Protein-containing fractions were pooled and concentrated using 3 kDa MWCO concentrators (#MAP003C37, Pall). Protein was dialysed using snakeskin dialysis tubing (cat no. 10005743, Thermo Scientific) into TEV cleavage buffer (50 mM Tris-base, 0.5 mM EDTA, 150 mM NaCl, 1 mM DTT, pH 8) and 10 µl of in-house TEV protease [40] per 1 mg of protein was added and incubated overnight at 4 °C. His-tagged TEV was separated from cleaved proteins using a Ni-NTA spin kit (#31314, Qiagen), and fractions were analysed by SDS-PAGE. Fractions containing cleaved protein were concentrated and further purified via size exclusion chromatography (Superdex75, GE Healthcare) using 50 mM Tris-base, 150 mM NaCl, pH 7.5. Protein-containing fractions were concentrated up to 14 mg/ml. The same purification procedure was employed for perdeuterated protein, with an additional dialysis into Tris/NaCl buffer in D_2_O for final H/D exchange of the labile hydrogens.

#### FtsH-IMS

FtsH-IMS was purified from lysate after high-speed centrifugation to remove unbroken cells using immobilised metal chromatography (IMAC) and anion exchange chromatography (AEC). IMAC was carried out using Ni-NTA Sepharose™ 6 Fast Flow resin (QIAGEN Ltd.) by first equilibrating it with 20 mM Tris, pH 7.5, 100 mM NaCl. The cell paste was sonicated and spun at 16,000 g for 60 minutes. The supernatant was diluted 1:3 with 20 mM Tris, pH 7.5, 100 mM NaCl and applied to the equilibrated column. The Ni-NTA resin was eluted with a step gradient of 20, 50, 100, 150, and 250 mM imidazole in 20 mM Tris, pH 7.5, 100 mM NaCl. Fractions were collected and proteins were detected using SDS-PAGE and protein-containing fractions pooled. AEC was performed using Q-Sepharose (Amersham Biosciences, GE Healthcare UK Ltd) employing a gradient from 0 M to 1 M NaCl in 20 mM Tris, pH 9.0. Fractions were analysed on SDS-PAGE and pooled before concentrating using a 3 kDa MWCO concentrator (Pall).

### 2.5 Mass spectrometry

FtsH-IMS was validated using mass spectrometry (MALDI-MS) carried out at the COIL facility, University of Edinburgh, Scotland, UK. Both intact protein mass determination and peptide-fingerprinting were performed.

For molecular mass determination of purified proteins, 3 µl of a 10 mg/ml protein solution was mixed with 27 µl of 50 mM ammonium bicarbonate, and 0.5 µl spotted onto the plate followed by overlay with 0.5 µl of sinapinic acid. Samples were then analysed by MALDI-MS.

For peptide fingerprint analysis, proteins were digested with porcine trypsin (Roche) to generate peptides for identification, either in-solution or in-gel according to sample quality. For in-gel digestion, SDS-PAGE gels were, stained with GelCode Blue Stain Reagent (Thermo Scientific) and destained with water. Protein bands were excised and SDS removed via washing 3 times with 300 µl buffer 1 (200 mM ammonium bicarbonate in 50 % acetonitrile) with samples left to incubate at room temperature for 30 minutes during each wash step. The gel was incubated in 300 µl buffer 2 (200 mM DTT, 200 mM ammonium bicarbonate, 50% acetonitrile) for one hour to reduce the protein and again washed three times with 300 µl buffer 1 before alkylation of cysteine residues in 100 µl buffer 3 (50 mM iodoacetamide, 200 mM ammonium bicarbonate in 50 % acetonitrile) at room temperature, dark conditions for 20 minutes. Gel pieces were washed three more times with 500 µl buffer 4 (20 mM ammonium bicarbonate in 50 % acetonitrile) and cut into 2 mm x 1 mm pieces. Samples were centrifuged (13000 g and 4 °C), supernatant discarded, and gel pieces covered with 100% acetonitrile for dehydration. Acetonitrile was removed, and dehydrated gel pieces allowed to dry. Gel pieces were transferred to chilled trypsin solution (1 µl trypsin stock solution and 29 µl 50 mM ammonium bicarbonate) and digested at 32 °C for 22 hours. For in-solution digestion, protein was denatured by first adding 8 M urea to give a protein concentration of 10 µg/µl, then 45 mM DTT was added, mixed and heated to 50 °C for 15 minutes. After cooling to room temperature 5 µl of 100 mM iodoacetamide was added and the sample incubated for 15 minutes at room temperature. A 20 µl aliquot of water was added to dilute the urea concentration to 2 M before adding 4 µl of trypsin stock solution and incubation at 27 to 28 °C for 18 to 24 hours. The reaction was stopped by freezing.

Following digestion, DTT was added to a final concentration of 5 mM. Samples were heated to 60 °C for 30 minutes, iodoacetamide added to a final concentration of 15 mM, and samples incubated for 30 minutes in the dark at room temperature. Samples were then sonicated prior to spotting onto the plate. For analysis by mass spectrometry, 0.5 µl of sample and 0.5 µl of α-cyano-4-hydroxycinnamic acid (CHCA) were spotted on a gold plate, left to dry before insertion into the mass spectrometer. Peptide fingerprints obtained were analysed with Data Explorer [41], and peaks analysed against the Mascot database [42] or the peptide database SwissProt (www.expasy.org). Results were analysed using the probability based Mowse score, −10 × log(*P*), where *P* is the probability that the observed match is a random event. Scores greater than 67 are considered significant (p < 0.05).

### 2.6 Circular dichroism (CD)

#### FtsH-IMS

CD cuvette with a light path of 1 mm (Hellma GmbH & Co. KG, Germany) and a Jasco J-810 spectropolarimeter equipped with a Peltier element were used to record wavelength spectra from 260 nm to 190 nm at 20 °C. The spectra were deconvoluted using the programme ACDP [43] to calculate mean residue ellipticity, Θ_mean,_ and implementations of two algorithms, K2D Neuronal Network and Linear Combination of Fasman Spectra were utilised for secondary structure deconvolution. Thermal stability of the protein was investigated by recording the change in the CD signal at 222 nm with a temperature slope of 1 °C /minute during heating of the sample from 20 °C to 80 °C. Changes in the CD signal were used to construct an unfolding curve which was fitted with SigmaPlot [44] using a sigmoidal equation. CD signals of His_6_-FtsH-IMS were monitored at a protein concentration of 0.04 mg/ml in 5 mM Hepes, pH 7.5, 20 mM NaCl.

#### Paraplegin-IMS

Circular dichroism (CD) was conducted on a Chirascan V100 instrument equipped with a Peltier temperature controller. CD spectra were collected from 180-280 nm using a cuvette with 1 mm pathlength (# 110-1-40, Hellma). For wavelength spectra acquisition, fixed interval scanning was performed at every nm, at 50nm/min scan speed with 5 data acquisitions, averaged. CD data in the range of 180-250 nm was deconvoluted using BeStSel [45]. Wavelength temperature interval data was collected between 25 to 90 °C every 2 °C, at 222 nm.

### 2.7 Small-Angle Neutron Scattering (SANS)

Small-Angle Neutron Scattering (SANS) data were acquired on D11 [46] at the Institut Laue-Langevin (ILL), Grenoble, France (D11_ILL_exp_8-03-946_201810132). An 3He MWPC detector of 0.96 × 0.96 m² was used, with 256 × 256 pixels. Samples were placed in quartz cells (Hellma, 110-QS) of 1 mm pathway, with an illuminated cross-section of 7 × 10 mm². A wavelength λ of 5.5 Å (relative FWHM 10 %) was selected, and data were recorded at 3 configurations, with sample-to-detector distances of 1.4, 5.5 and 8 m (collimation at 4, 5.5 and 8 m respectively), covering a *q*-range of 0.009–0.42 Å^-1^, where *q* is the magnitude of the wave-vector (*q* = 4π/λ sin(θ/2), θ being the scattering angle). For selected samples, additional measurements were performed at the longest detector distance (39 m) to reach lower *q* in order to confirm the intensity plateau obtained at higher *q*-values. Data were corrected for the relative pixel efficiencies, detector noise and dead-time, transmission, and incoming flux. The contribution of the empty cell was subtracted, and absolute scale was obtained using the intensity of H_2_O (1 mm pathway) as a secondary standard. 2D data were finally azimuthally averaged. Data are available on demand (DOI: 10.5291/ILL-DATA.8-03-946) and at DOI: 10.13140/RG.2.2.31690.27846.

### 2.8 Small Angle X-ray Scattering (SAXS)

#### Para-IMS

SAXS was performed at the BM29 beamline (European Synchrotron Radiation Facility, Grenoble, France, MX-1987, ESRF-BM29-20180508) [47]. BM29 was equipped with equipped with PILATUS 1 M detector (DECTRIS Ltd., Baden-Daettwil, Switzerland), the wavelength (λ) of the beam was 1.00 Å, giving *q* values ranging from 0.004 to 0.493 Å^−1^. To check for concentration effects, immediately before bioSAXS analysis samples were subject to a dilution series halving their concentration up to 1/8^th^ dilution with dialysis buffer. Initial concentrations of between 2-8 mg/ml were used. SAXS data was collected as 10 x 1 s exposure on 100 µl samples at 25 °C. A continuous flow cell capillary was used to reduce radiation damage. The data was averaged, the frames compared, and those displaying significant alterations discarded. SAXS profiles were subtracted by the corresponding blanks. ATSAS package [48] was used to analyse the data, with GNOM [49] used to estimate the radius of gyration (*R_g_*) in the Guinier region where *qR_g_* < 1.3. Plots were generated using GraphPad Prism (Dotmatics). Data are available on demand at DOI: 10.13140/RG.2.2.3001686 and DOI: <u>10.15151/ESRF-ES-118447013.</u>

#### FtsH-IMS

SAXS data were collected using the flux- and background-optimized NanoSTAR pin-hole camera (Bruker AXS) developed at Aarhus University [50]. The instrument uses Cu rotating anode in combination with Göbel mirrors and a HiSTAR 2D detector (Bruker AXS). Protein was measured at concentrations of 10 mg/ml, 5 mg/ml, and 2.5 mg/ml. Scattering data were corrected for background and converted to absolute scale using water as standard [50]. Based on its pair distance distribution function, *p*(*r*), obtained from Indirect Fourier transformation [51, 52], the curve for 5 mg/ml was selected for modelling the dimer. The modelling of the SAXS data was based on optimizing predicted 3D coordinate models until their theoretical scattering matches the experimental data following the procedure described in [53]. The structure is divided into several bodies and connectivity and excluded volume constraints are applied to get physically reasonable models. The models are optimized by random Monte Carlo moves, which are gradually reduced in size during the minimization of the agreement with the experimental SAXS data, which is expressed as the reduced chi-squared (χ^2^). At each step, a hydration layer is included to account for the ordered first layer of water around the protein. Ten runs were made for each data set to assess the variations in the models due to the random search procedure and the model for which the scattering gave the best fit to the data was selected as the resulting structure. Data are available on demand at DOI:10.13140/RG.2.2.16367.09121.

### 2.9 X-ray Crystallography

Crystallization experiments were carried out at the High Throughput Crystallisation Laboratory (HTX Lab) of the EMBL Grenoble, and crystals prepared for X-ray diffraction experiments using the CrystalDirect technology [54, 55]. Hydrogenated, TEV-digested para-IMS at 9.2 mg/ml was used to set up sparse-matrix screens in 96-well format, sitting drop, and small crystals grew in 0.17 M ammonium sulfate, 25.5% PEG 4000, 15% v/v glycerol. Larger crystals were grown after optimisation in 24-well plates using 1 µl 100 mM citrate buffer pH 4.2 with 10% PEG 8000, 1µl protein at 14.4 mg/ml and 0.5 µl seed stock prepared from the initial hit (1:10 diluted with crystallisation buffer) and hanging drop method. Crystals grew to around 0.1 mm x 0.13 mm. Dataset was collected at Diamond, 2000 images at 0.1° angle and 198 mm distance (BAG mx19880-27, beam line I04, wavelength 0.979515Å).

Indexing, integration and scaling were performed with the Xia2-dials workflow [56, 57] with automated indexing, refinement and integration, followed by pointless [58] and AIMLESS [59]. Molecular replacement was performed using Phaser [60] with a single chain model of the protein sequence generated via Alphafold3 [61] as template. For structure refinement and modelling, the programs Refmac [62] and Coot [63] were used, and Procheck [64] and Molprobity [64] for structure validation, as implemented in the CCP4 package. Data were deposited to the Protein data bank under PDB code 9RK6.

## 3 Results

### 3.1 The N-terminal regions of AAA-proteases are structurally conserved

Primary and secondary structure elements for the N-terminal regions of human paraplegin and AfG3L2 and bacterial FtsH were compared to identify conserved structural motifs which may pertain to function. For primary structure evaluation, pairwise sequence alignment of the N-terminal IMS regions of human paraplegin (residues 163 to 249), FtsH (residues 20 to 97) and human AfG3L2 (residues 168 to 248) revealed relatively low sequence similarity between Para-IMS and FtsH-IMS (14%), and Para-IMS and AfG3L2-IMS (34%). Despite poor alignment of primary sequences, secondary structure prediction revealed generally conserved structural elements across proteins: two hydrophobic transmembrane α-helices which flank an IMS region with well-defined secondary and tertiary structure elements. Specifically, the N-terminus is comprised of a hydrophobic α-helix (TM1, residues 3-21 - FtsH, 145-165 – SPG7, 143-163 - AfG3L2), followed by the IMS region, comprised of an α-helix, three β-strands, β1 to β3, another two short α-helices, α1 and α2, which can also present as one helix, and one short β-strand β4, which connects to a second hydrophobic α-helix (TM2, residues 97-121 - FtsH, 249-269 - SPG7, 251-271 – AfG3L2), bridging the IMS region to the catalytic ATPase and protease domains (Supplementary Figure S 1), also reported in [31]. The first and last helices of the N-terminus are believed to be transmembrane helices owing to their length relative to the thickness of the mitochondrial membrane and general hydrophobic character. In order to strengthen this hypothesis, the topology of FtsH as an integral membrane protein was predicted using MEMSAT [65]. It revealed two possible transmembrane helices, one from amino acids 2 to 20 (in ➔ out), the other spanning amino acids 97 to 124 (out ➔ in), and both helices were annotated TM1 and TM2, respectively. Therefore, the region between those two transmembrane helices represents an autonomous folding unit, with a defined 3D structure that carries a distinct function in AAA proteases. To investigate this further, we set out to produce the recombinant proteins of the human paraplegin and bacterial FtsH encompassing these regions, termed paraplegin-IMS and FtsH-IMS, respectively, and examine their molecular structure and oligomerisation states.

### 3.2 Recombinant paraplegin-IMS is a monomeric protein whereas FtsH-IMS appears as a dimer in solution

We successfully expressed paraplegin-IMS as a recombinant, His6-tagged protein from *E. coli* to a high degree of purity using nickel affinity chromatography at the expected size of 14.5 kDa (Figure 1A). Expression yields of 0.5 mg of purified and cleaved hydrogenated Paraplegin-IMS were achieved per litre bacterial culture, and ∼1 mg per 1 g cell paste for deuterated protein. Monodispersity of the fully-cleaved protein was indicated by a single peak at an apparent molecular weight of around 10 kDa in size exclusion chromatography which is in line with the calculated molecular weight of 9141 g/mol, whereas uncleaved protein has a calculated molecular weight of 14,578 g/mol (Figure 1A,B).

**Figure 1.**
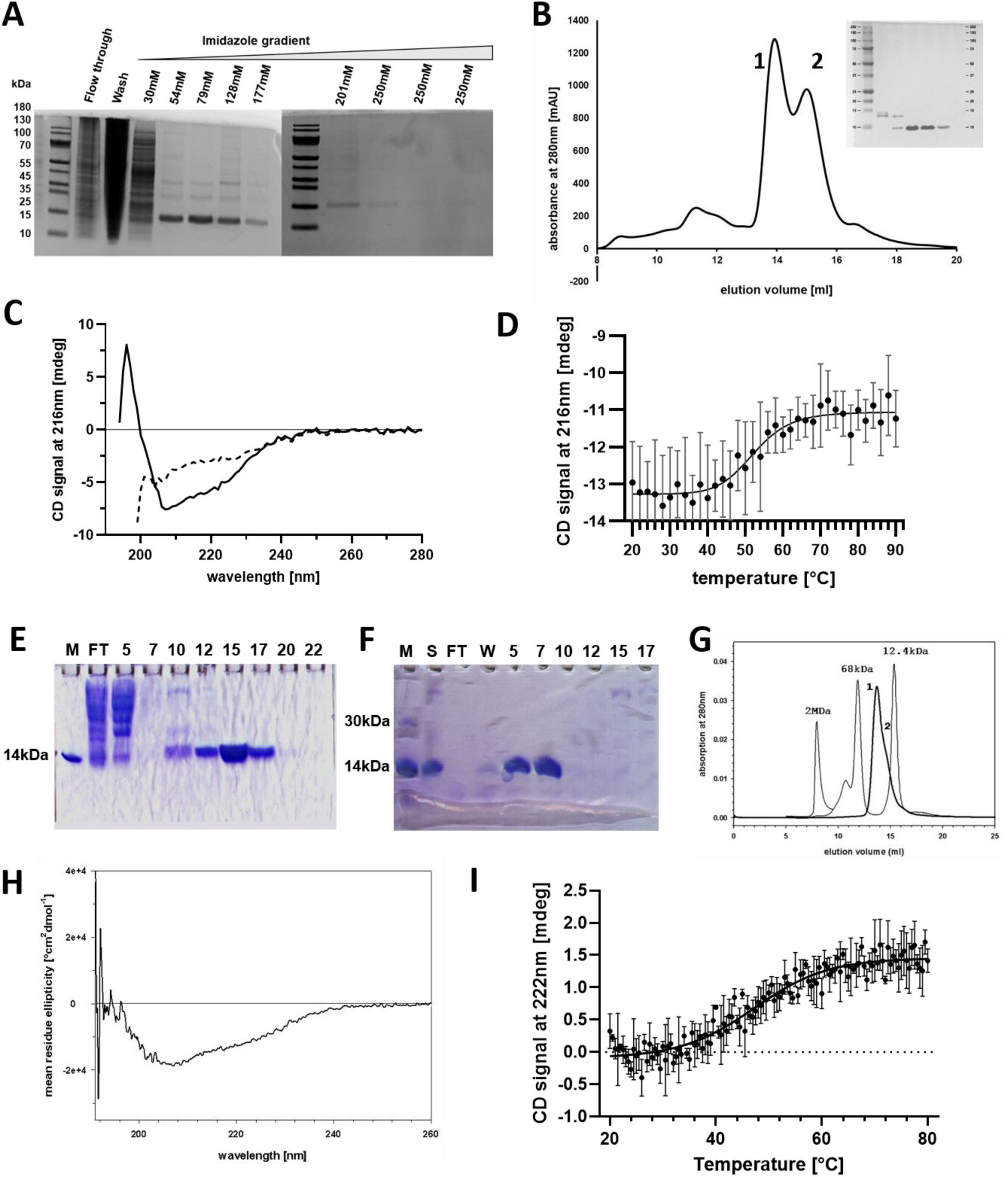
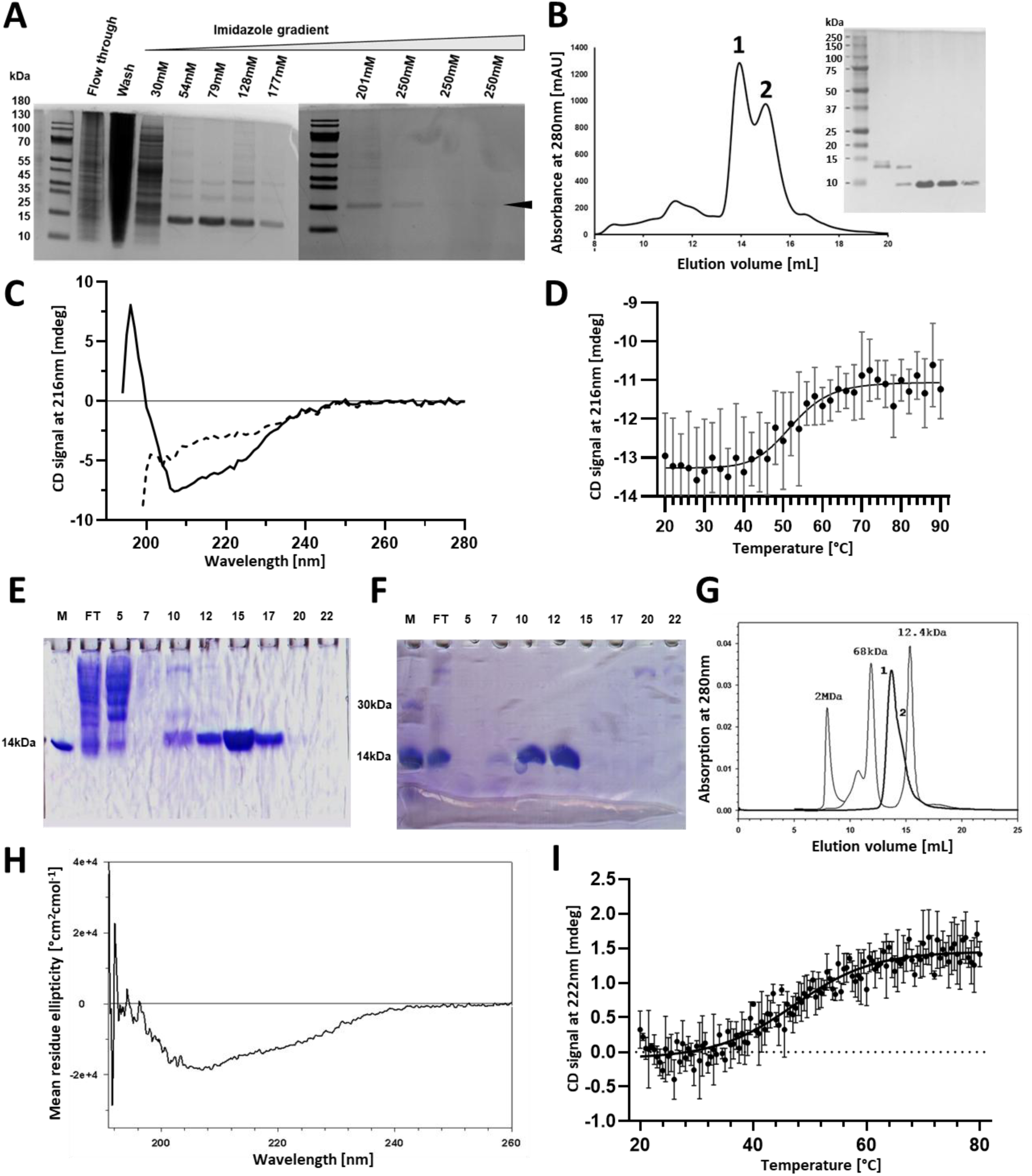
Fold and stability of recombinant paraplegin-IMS and FtsH-IMS. (A) IMAC purification of refolded, un-cleaved Para-IMS, showing on SDS-PAGE flow through (FT), wash (W), elution fractions of increasing imidazole concentration. His6-tagged Para-IMS is at an approximate size of 14k Da. (B) Size exclusion chromatography and accompanying SDS-PAGE after TEV-cleavage showing His6-Para-IMS as peak 1, just below 15 kDa on the SDS-PAGE, and cleaved Para-IMS (peak 2) at 10 kDa. Both hydrogenated and deuterated proteins showed similar profiles in IMAC and gel filtration. (C) CD wavelength spectrum at 20 °C (solid line) and 90 °C (dashed line) confirming a folded paraplegin-IMS protein. (D) Thermal denaturation of paraplegin-IMS protein and sigmoidal fit to determine the melting temperature *T_m_* at 208 nm with 48 °C. Shown are averages and standard deviations for three experiments. (E) SDS-PAGE of IMAC purification of His_6_-FtsH (20-97) with reference protein lysozyme in lane 1 (M), flow through (FT) and elution fractions in following lanes labelled by their fraction numbers. (F) SDS-PAGE from anion exchange chromatography with the reference proteins lysozyme and annexin A1 in lane 1 (M), sample (S), flow through (FT) and wash (W) and elution fractions labelled by their fraction numbers in following lanes. (G) Size exclusion chromatography in conjunction with light-scattering analysis reveal a peak 1 at around 26 kDa to 30 kDa with a shoulder as peak 2 around 15 kDa. Elution profiles of known gel filtration standard proteins (BioRad) known marker proteins shown for comparison. (H) CD wavelength spectrum of FtsH-IMS showing folded protein. (I) Thermal denaturation of FtsH-IMS region at 222 nm displays a two-state transition curve with a melting temperature *T_m_* of 49 °C. Shown is the average of three independent experiments and their standard deviation as error bars with fit as solid line.

FtsH could also be expressed as a His6-tagged protein to high yield and purified to purity, with a clear band visible on SDS-PAGE at ∼14 kDa (Figure 1E, F). The mass spectrum (MALDI-MS) shows a single dominant peak at 13,471 g/mol which is within range of the expected 13,488 g/mol (Supplementary Figure S 1A) for His6-FtsH (20-97). The identity of FtsH was verified by trypsin digest and subsequent analysis by mass spectrometry (Supplementary Figure S 2) employing a custom search database which found that 64% (76 of 117 amino acids) of the peptides matched that of the protein. The oligomerisation state of FtsH-IMS region was investigated using gel filtration analysis, where absorption peaks were fitted using the programme PeakFit [32], and apparent molecular masses of the peak maxima determined to be 26 kDa (peak 1) and 15 kDa (peak 2) (Figure 1G). Results suggest that the FtsH-IMS region eluted as a mixture of dimers and monomers with dimers as the predominant species, which contrasts with paraplegin-IMS, which was found to be monomeric. This was confirmed using SEC-MALS, where the molecular weight was determined to be 29 kDa, the approximate size of an FtsH-IMS region dimer. Dimer formation was also observed in the presence of 2 mM EDTA indicating that the dimer is not formed by the His6-tag. However, the dimers could be disrupted in presence of 1 M NaCl, and FtsH-IMS region eluted as a monomeric protein (results not shown), indicating monomer interaction that can be disrupted by increased ionic strength.

To further characterise both recombinant proteins with regards to their folding and stability, we conducted experiments using CD and DSF. However, data obtained with DSF were inconclusive (data not shown), possibly due to the small size of the proteins, as suggested elsewhere [66]. The fold of both proteins was therefore investigated by CD and subsequent deconvolution using BeStSel [67] to extract secondary structure contents. For FtsH-IMS, deconvolution revealed 7% α-helix, 31% in a β-sheet and 50% to be disordered (Figure 1H, Table 1) which is in agreement with secondary structure observed from X-ray crystallography (PDB code 4M8A). CD wavelength scans for paraplegin-IMS also indicated folded protein (Figure 1C) and were deconvoluted using BeStSel [67] with an RMSD < 0.06. Paraplegin-IMS consists of 34% β-strand, 14% turn, 7% helical, and 44% disordered structural elements. The β-strand amount is largely in agreement with published structural data from related proteins, e.g. Afg3L2 (PDB code 2LNA) with 32% β-strand and FtsH (PDB code 4M8A), with 31%. However, α-helical content appears to be quite significantly higher in published structures with 25% for Afg3L2 and 39% for FtsH. This may suggest a degree of disorder or flexibility in paraplegin where parts of the helices may not be formed appropriately and may adopt alternative conformations.

**Table 1:** Secondary structure analysis of paraplegin-IMS and FtsH-IMS.

| protein | technique | α-helix (%) | Turn (%) | β-strand (%) | Other (%) |
| --- | --- | --- | --- | --- | --- |
| Paraplegin* | CD | 7.4 | 14 | 34 | 44.6 |
| FtsH ** | CD | 6.7 | 12.9 | 30.7 | 49.8 |
| FtsH (4M8A) | X-ray crystallography | 39 |  | 31 | 30 |
| FtsH (4M8A) | CD prediction<sup>#</sup> | 26.6 |  | 33 |  |
| Afg3L2 (2LNA) | NMR | 25 |  | 32 | 45 |
| Afg3L2 (2LNA) | CD prediction<sup>#</sup> | 21.2 |  | 22.2 |  |

\* BestSel deconvolution with RMSD = 0.034
\*\* BestSel deconvolution with RMSD = 0.090

In summary, our results from CD experiments are largely in line with secondary structure assignments from published structural data, indicating a common secondary structure arrangement, as has been suggested previously [31, 32]. Of note is the assignment of some helical content to turns, which may indicate a degree of structural flexibility in solution. To further assess stability and fold of the proteins we conducted thermal denaturation experiments using CD. For paraplegin, sigmoidal fitting of data at 208 nm resulted in a two-state transition curve with a melting temperature *T_m_* of 48°C, indicating a stable, folded recombinant protein (Figure 1D). Unfolding of FtsH was monitored at 222 nm in three independent experiments and also shows a two-state transition curve with a melting temperature, *T_m_*, of 49 °C (Figure 1I). Our Results confirm that both paraplegin-IMS and FtsH-IMS contain a well-folded domain, and support the notion that the IMS-region of mAAA-proteases represents an autonomous folding unit with a mixed alpha-beta fold. We next set out to investigate their 3D structures *in solution* using SAXS and SANS.

### 3.3 *In-solution* small-angle scattering revealed a globular monomeric protein for paraplegin-IMS but distinct dimer formation for FtsH-IMS

Both experimental SANS and SAXS spectra of Para-IMS are consistent with a monomeric protein that contains a well folded, and globular domain (Figure 2A, B for SAXS and C, D for SANS). The radius of gyration (*R_g_*) is consistent for experimentally derived data from both SAXS and SANS at 14.3 Å and 12.3 Å, respectively. These values are in range of those obtained from simulated data of Afg3L2 NMR structure (PDB code 2LNA), using the PEPSI server [68], which give an *Rg* of 14.6 Å. This is further supported by Kratky analysis of the SAXS data which shows a typical profile for a well-folded globular protein (Figure 2C inset). Calculations of molecular weight return values of 7.3 kDa for SANS and 8.5 kDa for SAXS data, respectively, which is in good agreement with values obtained from size exclusion chromatography (∼10 kDa) and the theoretical value of 9.1 kDa for the TEV-cleaved protein. Pair distribution function *p*(*r*) analysis using GNOM [69] returned a maximum diameter of 35 Å for SANS and 40 Å from SAXS, respectively (Figure 2 B and D). Secondly, one representative of the NMR structure ensemble of Afg3L2 (PDB code 2LNA) was used as starting template to generate models that fit the SAXS data, albeit some low conservation in some regions (see Supplementary Figure S 1) and the presence of a purification tag. Most models have reduced chi-square (χ^2^) value around 1 (see Supplementary Table S 1), but models 3 and 8 give χ^2^ of nearly 3, and model 2 a χ^2^ of 2, with all of those models having parts sticking out, and therefore should probably be disregarded. The modelled structure of Para-IMS follows the same fold as observed in e.g. Afg3L2 NMR structure (PDB code 2LNA), but with some flexible extended loops and an extended N-terminal tail which represents the N-terminal His6-tag in the uncleaved protein (see Figure 2G). In conclusion, good models of paraplegin-IMS monomers could be obtained that fit our SAXS data well. Our data also show that paraplegin-IMS can be successfully produced in a deuterated form and that measurements in both SANS and SAXS reveal a globular monomer with approximate *R_g_* of 12 Å and 14.3 Å, respectively, and molecular weights of 7.3 kDa and 8.5 kDa, respectively. These experimental values are close to theoretical values calculated from its sequence.

**Figure 2.**
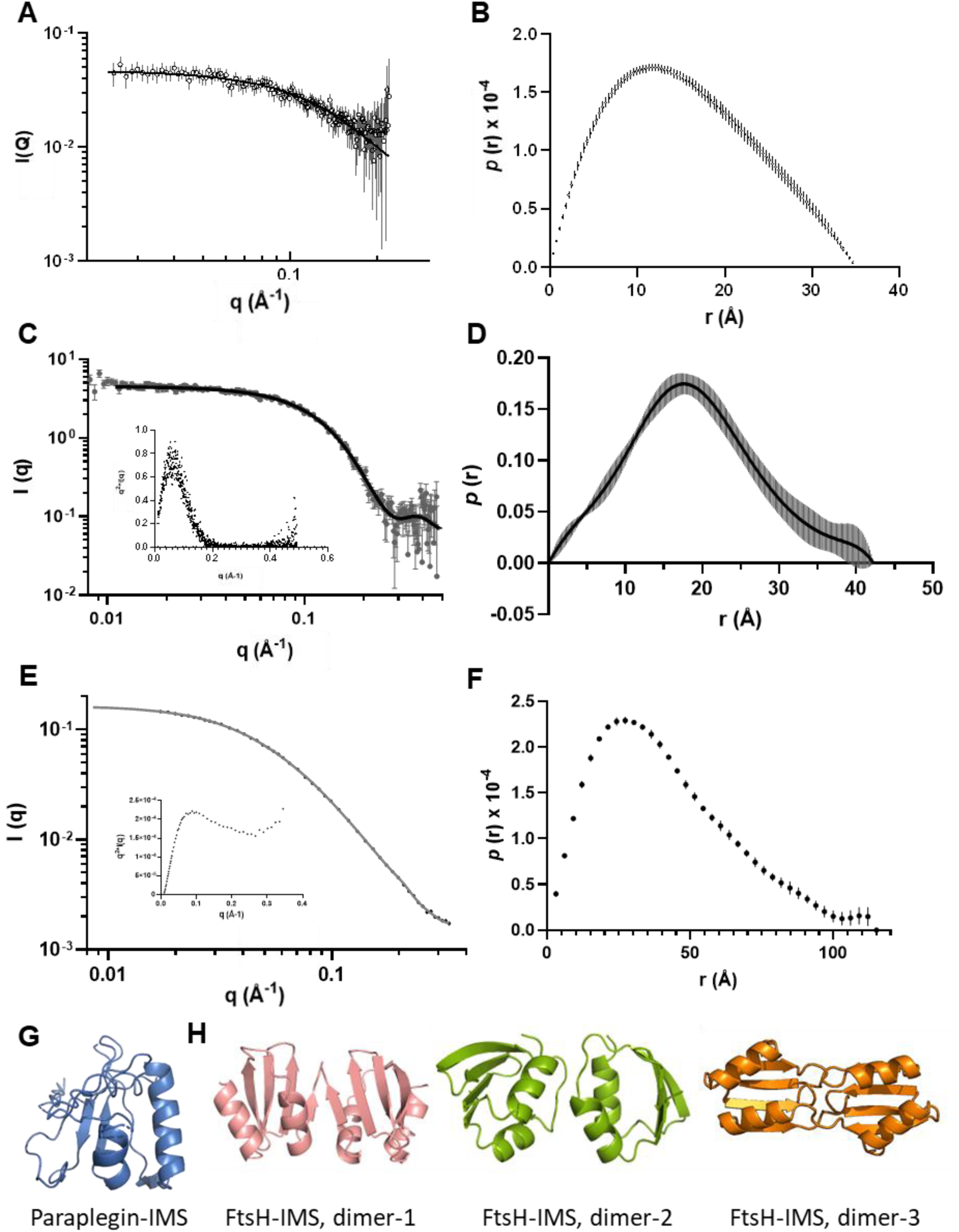
Analysis of SAS data for Paraplegin-IMS and FtsH-IMS. (A) SANS data of Paraplegin-IMS shown as circles with GNOM fit [69] shown as a line and corresponding *p*(*r*) distance distribution curve (B). (C) SAXS data of Paraplegin-IMS shown as circles with GNOM [69] fit shown as a line. Kratky plot as inset. (D) *p*(*r*) distance distribution function from Paraplegin-IMS SAXS data. (E) SAXS data of FtsH-IMS at 5mg/ml with model fit of dimer 3 shown as a line and Kratky plot as inset. (F) *p*(*r*) distance distribution curve from FtsH-IMS SAXS data. (G) Best model obtained for Para-IMS. (H) Best models obtained for putative FtsH dimers 1 (pink), dimer 2 (green) and dimer 3 (orange) omitting the flexible extended N-terminal tail containing the His6-tag.

The structural envelope of FtsH-IMS in solution was also examined using SAXS, with particular emphasis placed on identifying three-dimensional features relating to dimer formation (Figure 2E, F). To check for concentration effects, three concentrations of purified FtsH-IMS were analysed with SAXS: 10, 5 and 2.5 mg/ml. Some aggregation was observed, which complicated analysis of scattering data, but of the three concentrations measured, 5 mg/ml gave the best signal without artifacts, which after background subtraction and data normalisation was analysed with GNOM [69]. For all datasets, this *p*(*r*) analysis revealed a peak with shoulder, which is characteristic for dimers in solution (Figure 2F) and returned a maximum diameter of approximately 115 Å and *R_g_* of ≈33 Å, consistent with an elongated shape with a molecular mass of around 30 kDa as estimated from the forward scattering. Despite aggregation effects visible in the long tailing regions of the *p*(r) function, the data indicates a dimer, consistent with gel filtration analysis and in line with molecular weight estimated from SEC-MALS. A potential FtsH dimer in solution lacks direct evidence from cross-linking mass spectrometry; from both published data and our own attempts, which could support modelling and gain an understanding as to what the dimer interface could look like. Therefore, two different modelling approaches were employed, one using HADDOCK and one using AlphaFold. In the first case, two potential interaction sides were chosen based on available X-ray structures to generate two different dimers (dimer-1 and dimer-3). 3D coordinates of the FtsH monomer taken from structure 4V0B and the potential interaction interface was input into HADDOCK [70] using default settings, and 152 and 157 models were generated, respectively. Default clustering gave 18 and 9 clusters, respectively, from which two potential dimers with highest z-scores were obtained, dimer-1 (z-score of -1.7) and dimer-2 (z-score of -1.5), respectively (see Supplementary Figure 3). Fitting the dimer-1 model to the SAXS data gave eight models with χ^2^ values below 4, and five models with χ^2^ below 2 (Supplementary Table S 2). Fitting dimer-2 model to the data resulted in five models with χ^2^ below 2 (see Supplementary Table S 2). A third potential dimer was obtained by first generating models of an FtsH-IMS monomer from the primary sequence using AlphaFold and then using AlphaFold to generate a possible dimer. The putative dimer (dimer-3) was used as template to fit the SAXS data and 10 models were generated (see Supplementary Table S 2). Fits vary in χ^2^ values from 1-10, indicating that some models are a better fit to the SAXS data than others. Model 4 and 6 give the worst fits while model 2 gives the best fir with a χ^2^ values of <0.9. This analysis also indicates the presence of only a small fraction of dimers (10%) for the 5 mg/mL sample. Our analysis and simulations indicate that several dimer models could reproduce the experimental SAXS data, and we cannot conclude definitively whether one dimer arrangement is preferred over another, indicating a potentially quite flexible arrangement.

Dimer formation in IMS-regions of AAA-proteases may be linked to oligomer formation with implications for function, however, the arrangement at the molecular level is still unknown. To further investigate the tertiary structure and oligomeric organisation of these proteins, we proceeded to crystallise the IMS-region of paraplegin and to analyse its 3D structure at atomic detail using X-ray crystallography, which was compared to the previously published structures for FtsH from X-ray crystallography (4M8A), and for AfG3L2 from NMR (2LNA).

### 3.4 Crystal structure of paraplegin-IMS reveals a domain-swapped dimer

Initial crystallisation screens were set up at EMBL facility in Grenoble, France, and microcrystals obtained were used as seeds to further improve crystal size (Figure 3A). We obtained reasonably well diffracting crystals of TEV-cleaved Para-IMS (residues 163 to 249) to 2 Å in 2019 with unit cell parameters of a, b, c, as 56.39Å, 63.84 Å, 99.81 Å, space group C2 2 2_1,_ a water content of 45%, and likely two molecules in the asymmetric unit. We attempted to solve the dataset using molecular replacement and tools available at the time and with different modified and unmodified template structures of FtsH and AfG3L2 IMS-regions. However, we could initially not obtain a solution. When AlphaFold became available, the data was re-examined, and Alphafold [71] used to generate a template model which was less compact and in places more stretched than previously published experimental structures for FtsH and AfG3L2 IMS-proteins, which allowed us to solve the structure to 2 Å (Table 2).

**Figure 3.**
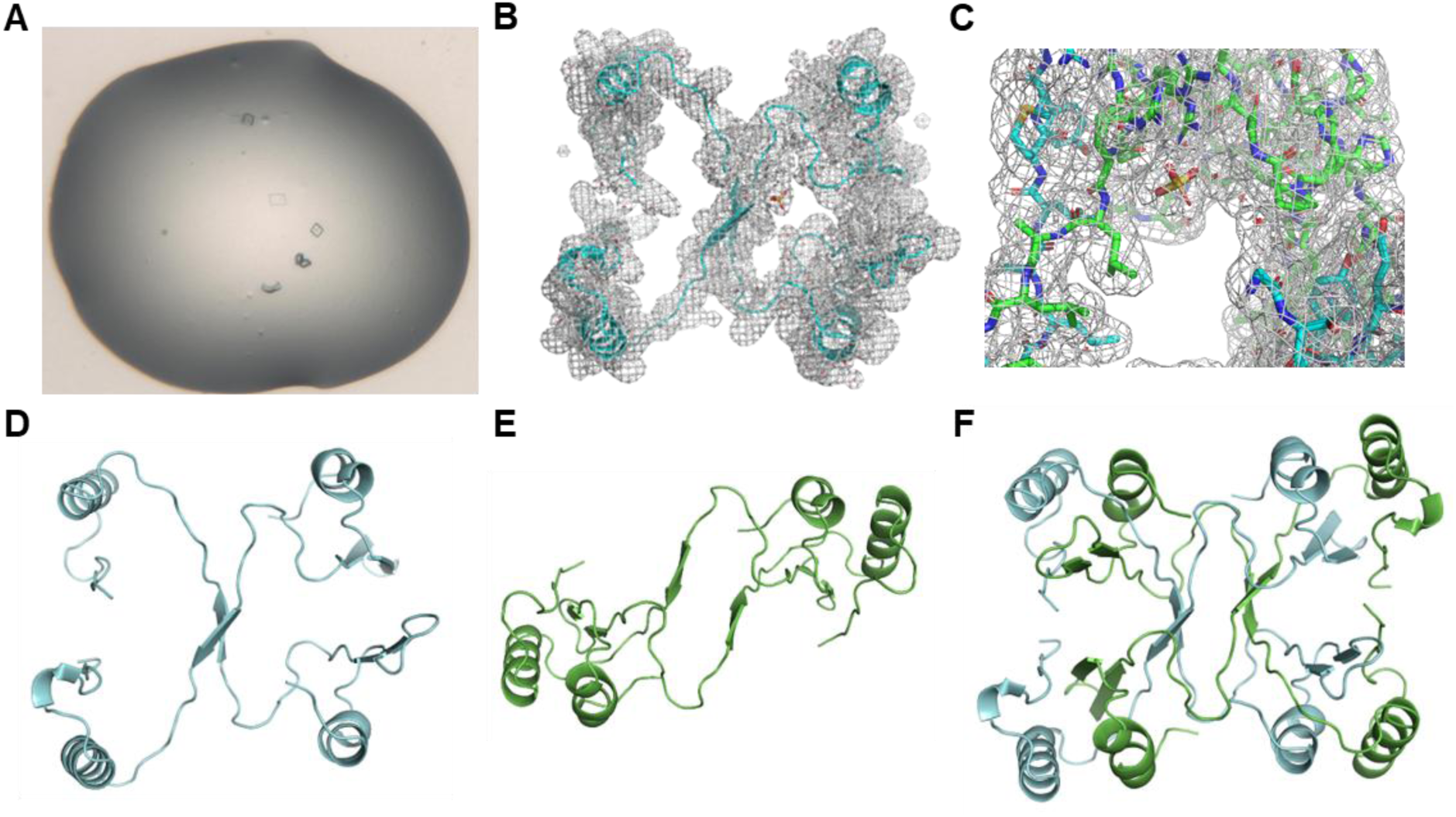
Domain-swapped crystal structure of paraplegin-IMS. A) Initial crystals from HTX screening. B - F) High-resolution structure of paraplegin-IMS in cartoon representation, (B) chain A and B of the asymmetric unit as the AB assembly with 2Fo-Fc map overlaid (grey), (C) Sulfate with an occupancy of 0.5 bound to AB assembly. Chain A shown in cyan and chain B shown in green (D) chain A and B of the asymmetric unit as the AB assembly (E) domain-swapped dimer of chain A with the appropriate symmetry-related molecule as the 2A assembly, (F) completed assembly 2A:2B of paraplegin-IMS with two molecules of one asymmetric unit shown in cyan and two symmetry-related molecules to complete the perceived monomer shown in green.

**Table 2:**
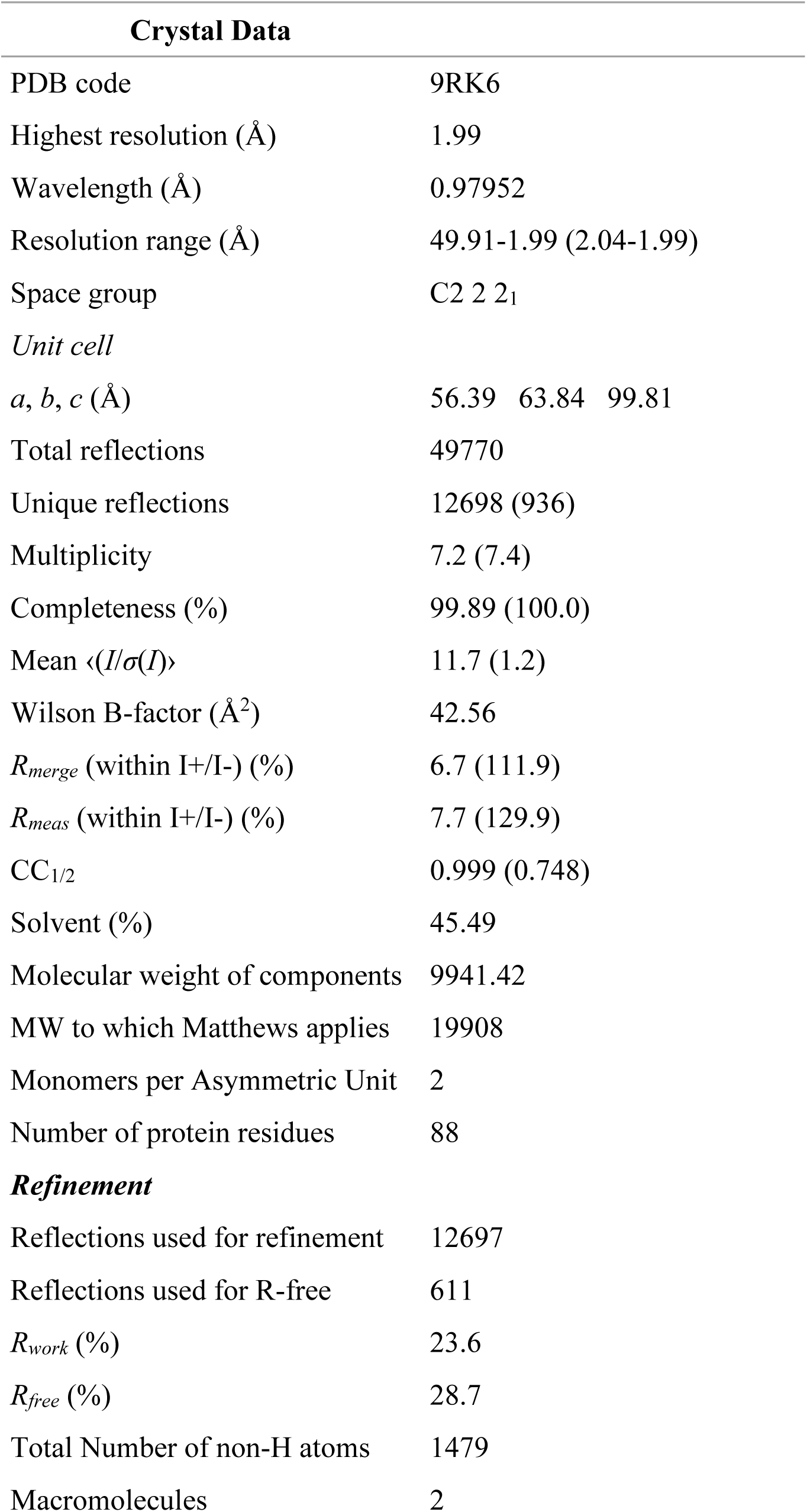
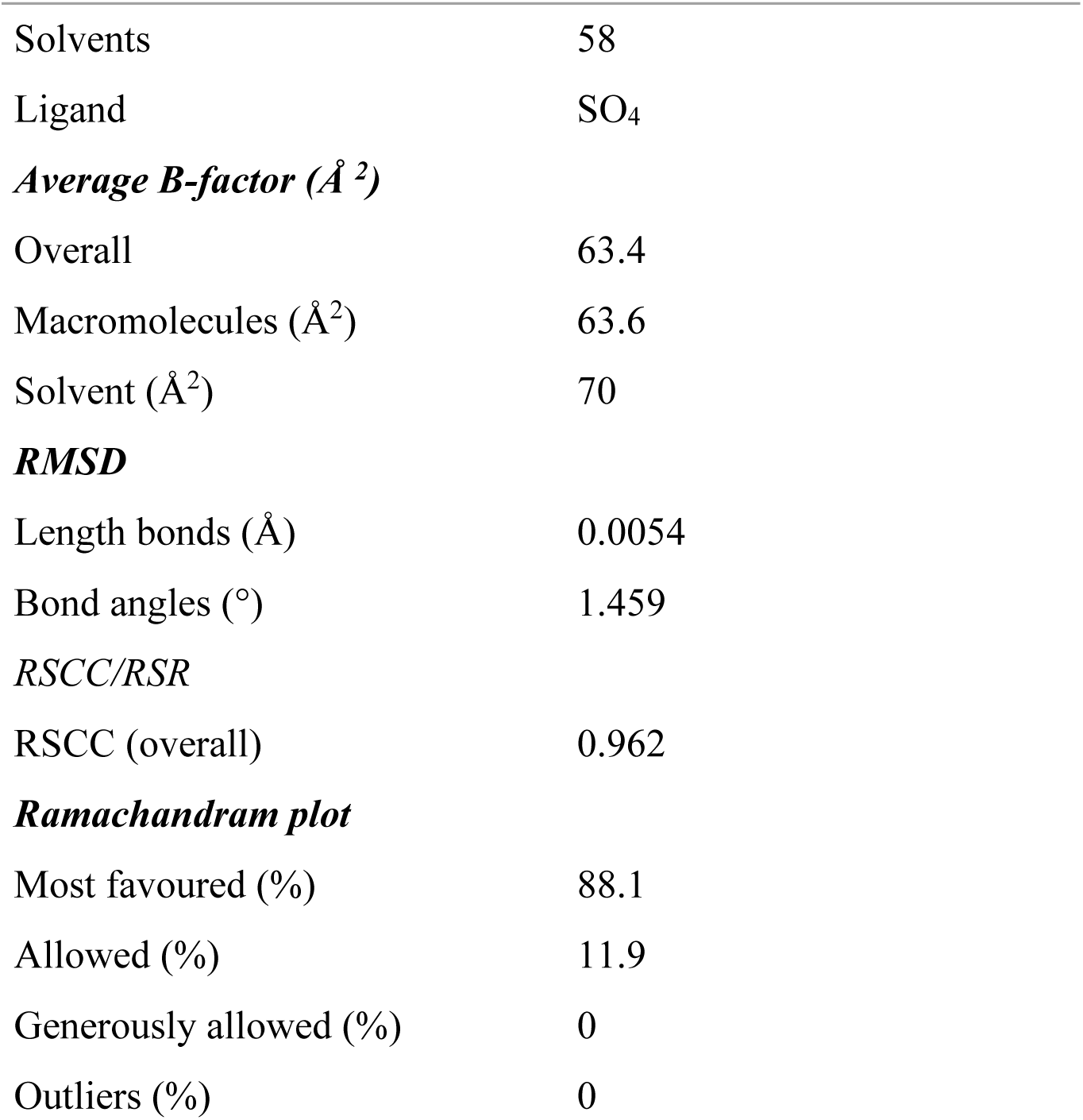
Data collection and refinement. Statistics for the highest-resolution shell are shown in parentheses.

This novel 3D structure revealed initially the same sequence of structural elements as observed for other N-terminal AAA-protease regions such as FtsH (PDB code 4M8A) and Afg3L2 (PDB code 2LNA); an α-helix, α1, followed by three β-strands, β1 to β3, then another α-helix, α2, one short β-strand, β4. The β-strands form a mixed beta-sheet, with β1 to β3 being antiparallel to each other whereas β4 is parallel to β3. The mixed sheet is flanked on one face by the two α-helices. However, two extended loops between β2 and β3 (12 residues) and between α2 and β4 (9 residues) were difficult to model in places due to a high degree of disorder and density being missing. This was especially true for the loop between β2 and β3 where density was missing between residues 46 and 47, and additional clear density was observed leading away from this connection. Closer examination of our data then revealed that the latter part of the protein was in fact donated by another molecule in the asymmetric unit, indicating domain swapping in our crystal structure. In essence, the N-terminal domain of paraplegin forms a domain-swapped homodimer that involves the first helix and first two beta-strands from one monomer and beta-strand 3, helix 2 and beta-strand 4 from another symmetry-related molecule (the final crystal data and refinement statistics are summarised in Table 2). This was very unexpected for such a small protein but could explain our difficulties in solving this dataset using molecular replacement with FtsH and Afg3L2 IMS-region structural data as templates. Trimming the data analysis using P1 space group essentially gave the same solution but with many more molecules in the asymmetric unit without improving refinement statistics, even after placing waters and other molecules. Re-examining the original images showed large and quite streaky spots which could be the reason for the higher than usual R factors and may point to a suboptimal crystal packing with some disorder. The crystals were quite small and fragile, and grew very slowly, which is potentially an indication of the disorder in the crystal packing. We hypothesise that a domain-swapped dimer is potentially a quite loose assembly that could form part of the active dimer in a hetero-hexameric complex with Afg3L2.

In detail, our crystal structure contains two molecules per asymmetric unit, A and B (AB assembly), each showing all structural elements apart from β4 which appears to be disordered. The structure is “ripped apart” between β2 and β3 with the latter part of the protein chain ∼34 Å away (Figure 3B). The two molecules interact chiefly via polar contacts from side chains, with two short beta-strands forming a central interaction. The interactions between the two molecules in the asymmetric unit are mediated by the following polar main contacts: A/Arg45 and B/Met49, A/Leu46 and B/Leu49 (main chain) and Arg45 (side chain), A/Ala47 and B/Ala47, A/Met49 and B/Arg45. In addition, two interaction points between side chains were identified: between Gln22, Tyr33, Arg20 of chain A with B/Gln19, and A/Gln19 with B/Tyr33. Interestingly, when chain A is shown with the appropriate symmetry-related chain A’ (2A assembly), the domain swapping becomes apparent, because their structural elements align at their N- and C-terminal ends to form what is the perceived monomer found in FtsH-IMS (e.g. PDB code 4V0B) or Afg3L2 (PDB code (2LNA) (Figure 3C and 4A, B). Both apparent monomers are separated by a long loop section that forms a beta-sheet half-way. Additional interactions are formed between α1, β1 and β2 of one monomer of the asymmetric unit and β2 and α2 of the symmetry-related monomer, with three beta-strands forming an anti-parallel sheet with a fourth disordered strand in some cases. The primary interface of the domain-swapping is composed of a large number of polar interactions between side chains, mainly facilitating the beta-sheet formation: A/Val21 with A’/ Pro80 and Val81, A/Val23 with A’/ Ser82 and A’/Lys 84, A/Asp28 and A’/Val54, A/Val30 and A’/Met52, A/Val32 and A’/ Tyr50, A/Try33 and A’/ Leu48. These are completed by polar side chain interactions between A/Ser2 and A’/ Gln53, and A/ Arg20 and A’/ Ser82. In addition to this, the symmetry-related molecules are interacting via a large network of hydrophobic residues consisting of: Trp5, Phe8, Val9, Leu13, Val30, Val21 and Val32 of the molecule in the asymmetric unit and Met52, Val54, Phe60, Leu64, Val81, Ala67, Leu71, Ile73 and Ile79 of the symmetry-related molecule. Intriguingly, the 2A assembly contains what appears to be a central oval cavity of around 25 Å by 7 Å which is flanked by two antiparallel beta-strands and one sulphate with 0.5 occupancy in each asymmetric unit. One could speculate that this channel might be quite flexible and allow a peptide chain to pass through and which might be involved in the peptidase activity of the ATP protease.

**Figure 4.**
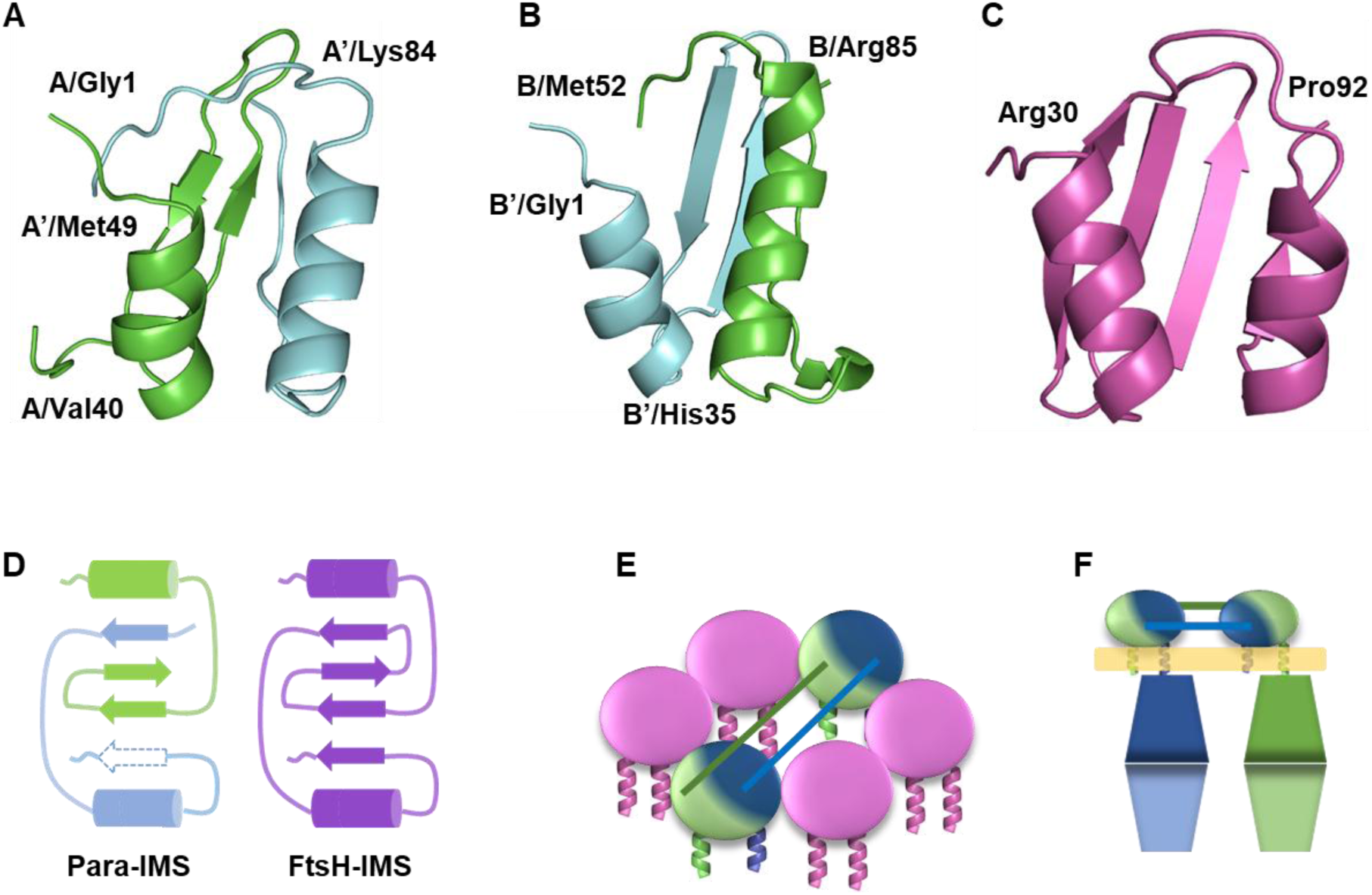
Proposed alternative oligomer formation in AAA proteases. (A) 2A dimer of paraplegin-IMS consisting of A/Gly1 to A/Val40 (green) from chain A and A’/Met49 to A’/Lys84 from the symmetry-related chain A’ (cyan). (B) 2B dimer consisting of B’/Gly1 to B’/His35 (cyan) from the symmetry-related chain B’ and B/Met25 to B/Arg85 (green) from chain B. (C) Arrangement of a FtsH-IMS protomer (PDB code 4V0B). (D) Cartoon schematic of structural arrangements in Para-IMS structure (left) with α1, β1 and β2 of one monomer of the asymmetric unit in green and β2 and α2 of the symmetry-related monomer in blue. The three beta-strands form an anti-parallel sheet with a fourth unordered strand indicated in dashes. Right: Comparison to FtsH structural arrangements with two helices and a central four-bladed mixed beta-sheet. (E) Proposed hetero-hexameric arrangement of m-AAA-protease consisting of paraplegin-IMS as a domain-swapped dimer (shown in green/blue) and four Afg3L2 protomers (magenta). (F) Proposed arrangement of paraplegin subunits in the hexamer including the matrix-side portions of the protein.

When examining an assembly of four chains including the appropriate symmetry-related protein chains A and B (2A:2B assembly) (Figure 3D) we notice that the beta-strands identified in the connecting extended loop sections of assemblies AB and 2A interact in such a way that they form opposing short anti-parallel beta-sheets which likely stabilise the tetramer assembly. However, we would like to point out that only the assembly of the domain-swapped chains A or chains B (2A assembly or 2B assembly) present with their N- and C-termini pointing to the same side. This is important because the IMS-region would be flanked by hydrophobic transmembrane helices, and only a correct arrangement of both the N-terminal and C-terminal ends of the IMS-region would therefore enable residence in the membrane. In contrast, the tetramer (2A:2B assembly) presents with opposite arrangements where the domain-swapped dimers are back-to-back. We therefore think it unlikely that the tetrameric 2A:2B assembly forms a valid biological assembly. We therefore propose that the 2A domain-swapped dimer is the biologically relevant arrangement whereas the 2A:2B assembly may be a crystallographic artefact.

## Discussion

Analysis of multi-domain membrane proteins represents a significant challenge in structural biology, especially for proteins which form large macro-molecular assemblies. Dissecting multi-domain membrane proteins into their constituent domains is a proven strategy for obtaining structural insights when full-length complexes resist conventional protein production procedures. Here, we present structural analysis of the N-terminal domain (IMS-region or external region or intermembrane domain) of the membrane-bound AAA proteases paraplegin and FtsH, whose function remains poorly defined, despite their apparent contributions to complex formation. We confirmed that these N-terminal domains present autonomous folding units, as has been suggested previously [28, 31], and were able to produce the fragments to high purity and yield using bacterial expression. Intriguingly, the IMS-region from human paraplegin presents as a stable monomer in solution and in contrast as a domain-swapped dimer when crystallised, whereas the IMS-region from bacterial FtsH presents mostly as a dimer in solution. This suggests an oligomer formation between N-terminal domain monomers that is independent of assembly of the hexameric C-terminal domain, and may therefore contribute to guiding oligomer formation, complex stabilisation or functionality of the complex.

Atomic-level structural information on the N-terminal IMS-region of AAA proteases exists from X-ray crystallography of bacterial FtsH from *Thermotoga maritima* (PDB codes 4M8A, 4Q0F) which presents as a dimer in the asymmetric unit. The protomers are structurally very similar to the X-ray structure of *E.coli* FtsH-IMS (4V0B, [32]) which presents as a hexamer in the asymmetric unit. A structurally similar monomer was obtained from NMR structures of *E.coli* FtsH-IMS (PDB code 2MUY, [32]) and human AfG3L2 (PDB code 2LNA, [33]). This apparent structural similarity between protomers from different orthologues suggests a common fold of this domain. We set out to determine the atomic structure of paraplegin-IMS and were able obtain crystals diffracting to 2Å. Surprisingly, we encountered difficulty solving the structure, which already pointed towards a non-canonical structural arrangement of the monomer units. We found that the asymmetric unit of paraplegin-IMS contains loosely interacting dimers which are further stabilised through domain-swapping interactions with symmetry-related dimers, in which structural elements are exchanged between neighbouring protomers (Figure 4A and B). The domain-swapped monomers are structurally similar to monomers obtained from previous structural studies (e.g. PDB codes 4M8A, 4Q0F, 2MUY, 2LNA) (Figure 4C), and structural overlays show that α1 and β1 to β3 generally align well while helix α2 is often slanted at an angle and β4 is not well-defined. Nonetheless, this points to a common fold of the N-terminal autonomous domains of AAA proteases across orthologues (Figure 4D).

In contrast, our solution scattering data indicate a monodisperse monomer, and recent cryo-EM structures of the homologous FtsH-HflK/C complex also lack evidence of domain swapping [78]. While this specific homodimer arrangement is possibly a crystallization artifact driven by non-physiological packing forces [79, 80], the crystal structure demonstrates that the paraplegin IMS domain can adopt this conformation, indicating an inherent capacity for conformational flexibility that may be functionally relevant in accommodating heteromeric assembly with Afg3L2 (Figure 4E,F).

It has previously been established that the IMS-region of AAA proteases facilitates complex formation of the functional, membrane-anchored, hexameric protein [28], and that removal of the IMS-region of FtsH leads to some loss of interaction within its homo-hexamer but also with important functional interaction partners, such as prohibitins [28, 72]. We did not observe the formation of a hexamer in solution, indicating that hexamer formation is either achieved via the C-terminal parts of the protein [17], mediated through binding partners [73] or by organisation within a lipid bilayer environment. This is supported by the cryo-electron structure of FtsH that shows a compact hexameric C-terminal ATPase and metallo-peptidase domain whereas the IMS-region is modelled as distinct protomers [17].

Comparison between the oligomeric states of the N-terminal regions of paraplegin and FtsH reveals markedly different features. Bacterial FtsH-IMS majorly forms homodimers under the conditions tested, whilst paraplegin-IMS can exist as stable monomers in solution or as domain-swapped homodimers as demonstrated by its crystal structure. This is likely indicative of differing requirements of each protomer at the hexameric level. In FtsH-IMS, the formation of a homodimer may serve as a nucleation point for hexamer formation through a “trimer of dimers” assembly, as has been proposed for related AAA+ systems [81]. In contrast, human paraplegin exhibits more varied assembly behaviour. Afg3L2 exists as homo-hexamers or forms hetero-hexamers with paraplegin, whilst paraplegin is only able to form hetero-hexamers with Afg3L2, allowing for a range of hexamer assemblies of varying composition (e.g. 2+4 or 4+2 stoichiometries) which may be related to specific functions [15, 76]. The extent of this variation across hexamers with regards to specific cellular localisation or cell types is poorly understood. However, biochemical and expression studies have shown tissue and cell-specific variation in paraplegin vs AFG3L2 abundance and assembly [15, 77]. Homo- and hetero-hexamers may confer distinct mechanistic properties [26] which may enable different functional requirements in specific cellular contexts arising from subunit composition.

We propose that the conformational flexibility demonstrated by paraplegin’s IMS domain — its capacity to unlatch and share structural elements — provides a structural basis for accommodating these varied heteromeric arrangements (Figure 4E). In our crystal structure, we identified an extensive network of polar and non-polar interactions capable of supporting exchange between neighbouring subunits via extended, flexible stretches. Domain-swapping is a well-documented mechanism for complementing prevailing secondary structural features [74, 75]. In addition to interactions formed primarily through the ATPase domain, this plasticity could support a stable hexamer while permitting necessary flexibility in the matrix-side domains (Figure 4F). However, while the IMS region may contribute to stability, substrate translocation and unfolding strictly require energy and must therefore remain functionally and mechanically coupled to the ATPase moiety.

The findings presented here confirm the basic structural assembly of FtsH-IMS and paraplegin-IMS, whilst highlighting their markedly different oligomeric behaviours in solution. While the specific homodimeric domain swap observed in our paraplegin crystal structure is not proven to occur physiologically, the underlying capacity for conformational flexibility is a bona fide structural feature of the protein. We therefore advocate for the re-examination of existing structural data for evidence of similar conformational plasticity, which may have been previously overlooked but could be essential for facilitating heteromeric assembly in this class of membrane proteins.

## Acknowledgements

This work was supported by access to the HTX lab facility at EMBL and the PSB. The authors would like to thank Jose A. Marquez and Guillaume Hoffmann for their support with using the facility. The research leading to these results has received funding from the European Community’s Seventh Framework Programme H2020 under iNEXT (grant agreement N°653706). We also acknowledge the ILL for access to the Deuteration Laboratory (ILL D-Lab) and provision of resources. The authors acknowledge support from beamline staff at I04, Diamond. We would like to acknowledge contributions from undergraduate student Yumna Ladha.

## Author contributions

JH: Data curation, Formal data analysis, Investigation, Visualization, Writing original draft, Review & editing

NGP: Data curation, Formal data analysis, Review & editing

JMD: Investigation, Review & editing

CLPO: Formal data analysis, Investigation, Review & editing

CMJ: Formal data analysis, Investigation

SP: Data curation, Formal data analysis, Investigation, Validation, Review & editing

JSP: Data curation, Formal data analysis, Investigation, Supervision; Validation, Review & editing

AH: Conceptualization, Supervision, Review & editing

AW: Conceptualization, Data curation, Formal data analysis, Investigation, Supervision; Validation; Visualization, Writing original draft, Review & editing;

## Supplementary Data

### Supplementary Figures

**Figure S1.**
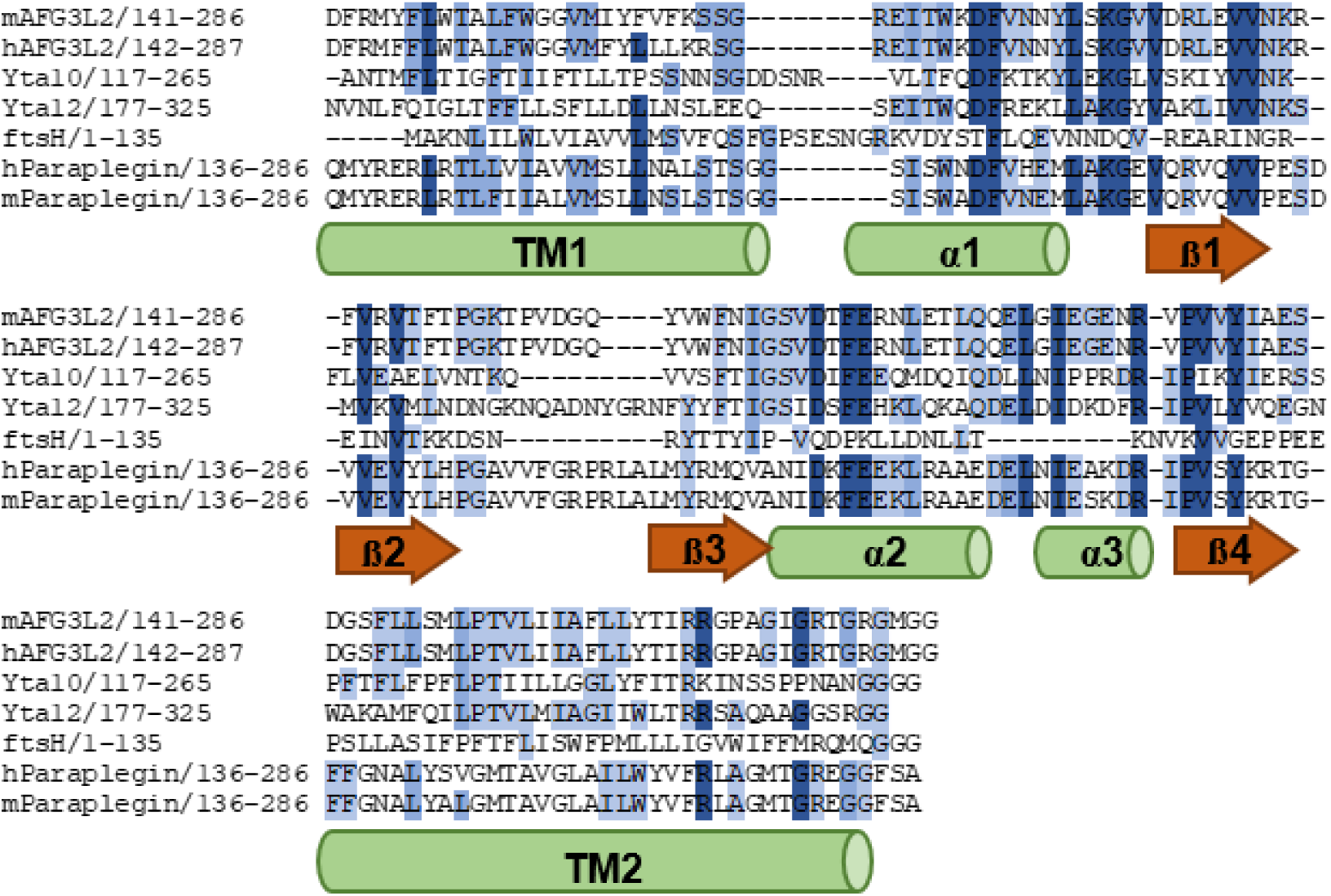
Structural conservation of the IMS region. Multiple sequence alignment of different AAA-protease subunits, Thermotoga maritima FtsH (UniProtKB ID P0AAI3, PDB code 4M8A), Yta10 (Sequence ID: X81066), Yta12 (UniProtKB ID: P40341), human AfG3L2 (UniProtKB (ID: Q9Y4W6), mouse AfG3L2 (UniProtKB (ID: Q8JZQ2, PDB code 2LNA), human paraplegin (UniProtKB ID: Q9UQ90), and mouse paraplegin (InterPro IPR005936) generated using, complemented with secondary structure predictions and annotations where available. Percentage identity indicated in shades of blue with > 40% light blue, > 60% medium blue and > 80% dark blue. The hydrophobic membrane-spanning helices TM1 and TM2 border a sequence of α-helices (green cylinders) and ß-strands (red arrows).

**Figure S2.**
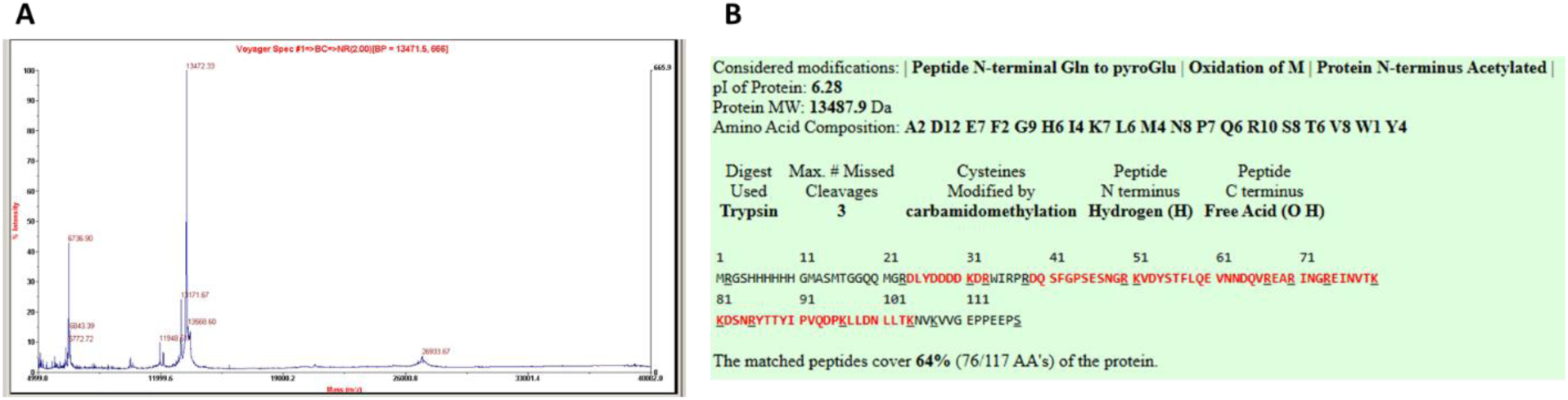
FtsH-IMS mass spectrometry results. A) MALDI-MS revealing a peak at 13472 g/mol B) finger print analysis revealing 64% sequence coverage.

**Figure S3.**
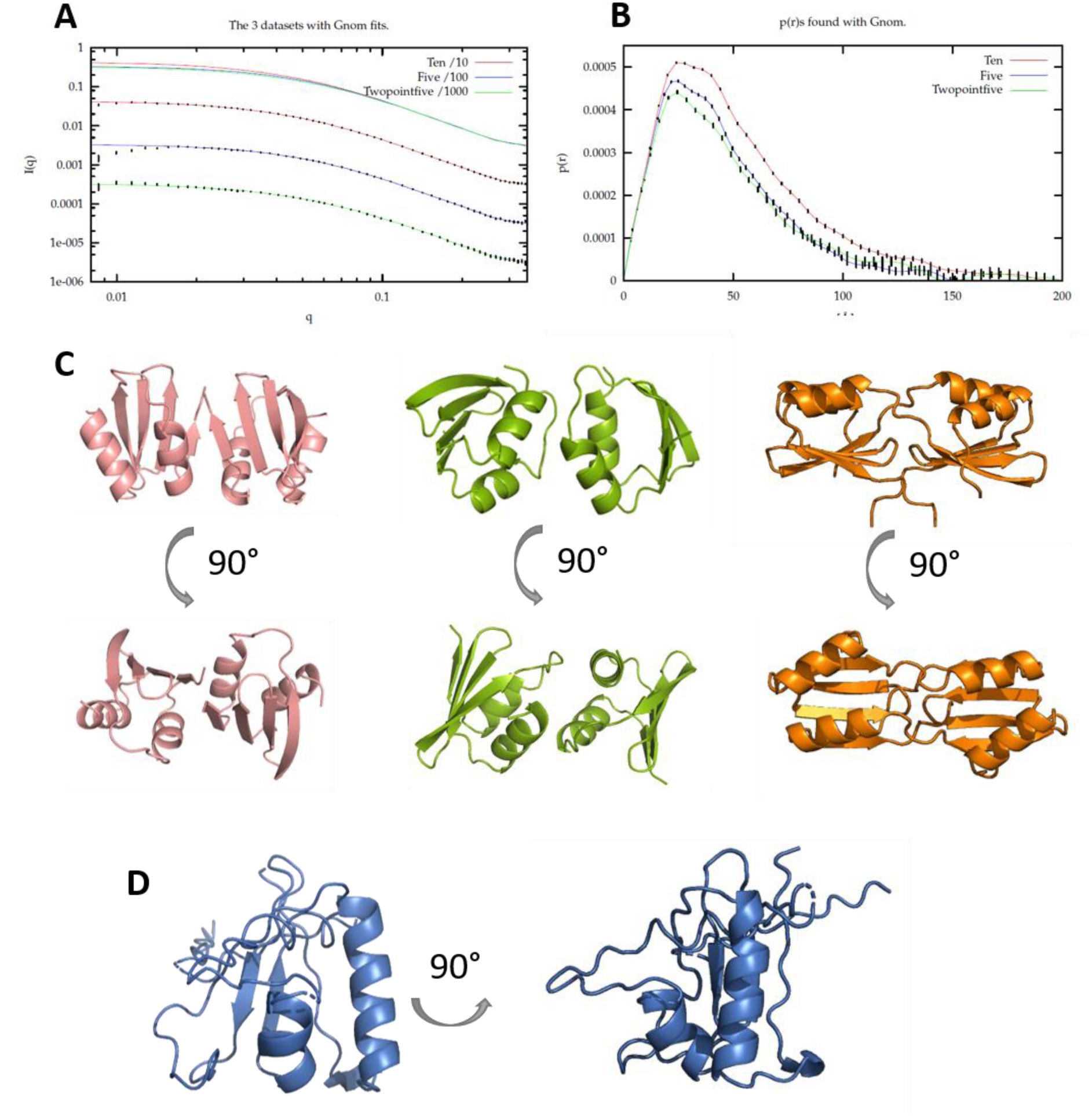
SAXS data of FtsH-IMS. A) SAXS data of FtsH at concentrations of 10 mg/ml (red), 5 mg/ml (blue) and 2.5 mg/ml (green). B) Corresponding pair distribution functions as determined using Gnom. C) Potential dimers formed by FtsH monomers in solution that could represent the SAXS profiles, modelled using HADDOCK shown as dimer-1 (magenta) and dimer-2 (green). Dimer model using AlphaFold (dimer-3) (orange). (D) The NMR structure of Afg3L2 (PDB code 2LNA). F) Afg3L2-IMS (PDB code 2LNA) (red) and proposed paraplegin-IMS swapped dimer (yellow and magenta) are overlaid for comparison.

### Supplementary tables

**Table S1.** Modelling of Para-IMS data using AlphaFold and subsequent rigid body refinement to create an initial model. . Model 6 is the best model.

| Model | $\chi^2$ |
| --- | --- |
| **1** | 1.21 |
| **2** | 1.98 |
| **3** | 2.87 |
| **4** | 1.39 |
| **5** | 1.41 |
| **6** | **1.21** |
| **7** | 1.23 |
| **8** | 2.86 |
| **9** | 1.22 |
| **10** | 1.24 |

**Table S2.** Modelling of FtsH-IMS SAXS data. . HADDOCK was used to create an initial dimer model (dimer-1, z-score of -1.7) with the best Monte Carlo model fit to the data for model 5. HADDOCK was used to create an initial dimer model (dimer-2, z-score of -1.5) with the best Monte Carlo model fit to the data for model 4. Reduced χ^2^ values of the Monte Carlo optimised fits to the data are shown for each model. AlphaFold-generated dimer (dimer-3) was used as template to fit the SAXS data, and model 2 represents the best fit.

| Model | Dimer-1<br>$\chi^2$ | Dimer-2<br>$\chi^2$ | Dimer-3<br>$\chi^2$ |
| --- | --- | --- | --- |
| 1 | 3.867 | 2.79 | 4.68 |
| 2 | 9.07 | 2.25 | **0.88** |
| 3 | 1.68 | 2.52 | 2.74 |
| 4 | 2.748 | **1.06** | 8.00 |
| 5 | **1.12** | 1.41 | 2.67 |
| 6 | 2.42 | 1.41 | 10.267 |
| 7 | 1.37 | 4.27 | 1.72 |
| 8 | 1.31 | 3.38 | 3.42 |
| 9 | 1.60 | 1.31 | 4.39 |
| 10 | 5.83 | 1.61 | 1.68 |

## References

1. Tomoyasu, T., et al., The Escherichia coli FtsH protein is a prokaryotic member of a protein family of putative ATPases involved in membrane functions, cell cycle control, and gene expression. J Bacteriol, 1993. 175(5): p. 1344–51.

2. Deuerling, E., et al., The ftsH gene of Bacillus subtilis is involved in major cellular processes such as sporulation, stress adaptation and secretion. Mol Microbiol, 1997. 23(5): p. 921–33.

3. Tatsuta, T., et al., m-AAA protease-driven membrane dislocation allows intramembrane cleavage by rhomboid in mitochondria. Embo J, 2007. 26(2): p. 325–35.

4. Kihara, A., Y. Akiyama, and K. Ito, Different pathways for protein degradation by the FtsH/HflKC membrane-embedded protease complex: an implication from the interference by a mutant form of a new substrate protein, YccA. J Mol Biol, 1998. 279(1): p. 175–88.

5. Kolodziejczak, M., R. Skibior-Blaszczyk, and H. Janska, m-AAA Complexes Are Not Crucial for the Survival of Arabidopsis Under Optimal Growth Conditions Despite Their Importance for Mitochondrial Translation. Plant Cell Physiol, 2018. 59(5): p. 1006–1016.

6. Pareek, G., R.E. Thomas, and L.J. Pallanck, Loss of the Drosophila m-AAA mitochondrial protease paraplegin results in mitochondrial dysfunction, shortened lifespan, and neuronal and muscular degeneration. Cell Death Dis, 2018. 9(3): p. 304.

7. Bulteau, A.L. and A. Bayot, Mitochondrial proteases and cancer. Biochim Biophys Acta, 2011. 1807(6): p. 595–601.

8. Rugarli, E.I. and T. Langer, Mitochondrial quality control: a matter of life and death for neurons. Embo j, 2012. 31(6): p. 1336–49.

9. Murru, S., et al., Astrocyte-specific deletion of the mitochondrial m-AAA protease reveals glial contribution to neurodegeneration. (1098-1136 (Electronic)).

10. Casari, G., et al., Spastic paraplegia and OXPHOS impairment caused by mutations in paraplegin, a nuclear-encoded mitochondrial metalloprotease. Cell, 1998. 93(6): p. 973–83.

11. Martinelli, P., et al., Genetic interaction between the m-AAA protease isoenzymes reveals novel roles in cerebellar degeneration. (1460-2083 (Electronic)).

12. Akiyama, Y., T. Yoshihisa, and K. Ito, FtsH, a membrane-bound ATPase, forms a complex in the cytoplasmic membrane of Escherichia coli. J Biol Chem, 1995. 270(40): p. 23485–90.

13. Krzywda, S., et al., The crystal structure of the AAA domain of the ATP-dependent protease FtsH of Escherichia coli at 1.5 A resolution. Structure, 2002. 10(8): p. 1073–83.

14. Koppen, M. and T. Langer, Protein degradation within mitochondria: versatile activities of AAA proteases and other peptidases. Crit Rev Biochem Mol Biol, 2007. 42(3): p. 221–42.

15. Koppen, M., et al., Variable and tissue-specific subunit composition of mitochondrial m-AAA protease complexes linked to hereditary spastic paraplegia. Mol Cell Biol, 2007. 27(2): p. 758–67.

16. Arlt, H., et al., The YTA10-12 complex, an AAA protease with chaperone-like activity in the inner membrane of mitochondria. Cell, 1996. 85(6): p. 875–85.

17. Lee, S., et al., Electron cryomicroscopy structure of a membrane-anchored mitochondrial AAA protease. J Biol Chem, 2011. 286(6): p. 4404–11.

18. Puchades, C., et al., Unique Structural Features of the Mitochondrial AAA+ Protease AFG3L2 Reveal the Molecular Basis for Activity in Health and Disease. Mol Cell, 2019. 75(5): p. 1073–1085.e6.

19. Steele, T.E. and S.E. Glynn, Mitochondrial AAA proteases: A stairway to degradation. Mitochondrion, 2019. 49: p. 121–127.

20. Piechota, J., et al., Identification and characterization of high molecular weight complexes formed by matrix AAA proteases and prohibitins in mitochondria of Arabidopsis thaliana. J Biol Chem, 2010. 285(17): p. 12512–21.

21. Vostrukhina, M., et al., The structure of Aquifex aeolicus FtsH in the ADP-bound state reveals a C2-symmetric hexamer. Acta Crystallogr D Biol Crystallogr, 2015. 71(Pt 6): p. 1307–18.

22. Langklotz, S., F. Baumann U Fau - Narberhaus, and F. Narberhaus, Structure and function of the bacterial AAA protease FtsH. (0006-3002 (Print)).

23. Langer, T., AAA proteases: cellular machines for degrading membrane proteins. Trends Biochem Sci, 2000. 25(5): p. 247–51.

24. Ogura, T. and A.J. Wilkinson, AAA+ superfamily ATPases: common structure--diverse function. Genes Cells, 2001. 6(7): p. 575–97.

25. Guha, S., et al., Transcriptional and cellular responses to defective mitochondrial proteolysis in fission yeast. (1089-8638 (Electronic)).

26. Bieniossek, C., B. Niederhauser, and U.M. Baumann, The crystal structure of apo-FtsH reveals domain movements necessary for substrate unfolding and translocation. Proc Natl Acad Sci U S A, 2009. 106(51): p. 21579–84.

27. Suno, R., et al., Structure of the whole cytosolic region of ATP-dependent protease FtsH. Mol Cell, 2006. 22(5): p. 575–85.

28. Akiyama, Y. and K. Ito, Roles of multimerization and membrane association in the proteolytic functions of FtsH (HflB). Embo j, 2000. 19(15): p. 3888–95.

29. Korbel, D., et al., Membrane protein turnover by the m-AAA protease in mitochondria depends on the transmembrane domains of its subunits. EMBO Rep, 2004. 5(7): p. 698–703.

30. Scharfenberg, F., et al., Structure and evolution of N-domains in AAA metalloproteases. (1089-8638 (Electronic)).

31. Winter, A., Structural Biology of Prohibitins and Annexin B1, in Dept of Biological Sciences. 2008, University of Edinburgh: Edinburgh.

32. Scharfenberg, F., et al., Structure and evolution of N-domains in AAA metalloproteases. J Mol Biol, 2015. 427(4): p. 910–923.

33. Ramelot, T.A., et al., NMR structure and MD simulations of the AAA protease intermembrane space domain indicates peripheral membrane localization within the hexaoligomer. FEBS Lett, 2013. 587(21): p. 3522–8.

34. Higgins D., T.J., Gibson T. Thompson J.D., Higgins D.G., Gibson T.J., CLUSTAL W: improving the sensitivity of progressivemultiple sequence alignment through sequence weighting,position-specific gap penalties and weight matrix choice. Nucleic Acids Res., 1994. 22: p. 4673–4680.

35. McGuffin, L.J., K. Bryson, and D.T. Jones, The PSIPRED protein structure prediction server. Bioinformatics, 2000. 16(4): p. 404–5.

36. Waterhouse, A.M., et al., Jalview Version 2--a multiple sequence alignment editor and analysis workbench. Bioinformatics, 2009. 25(9): p. 1189–91.

37. Studier, F.W., Protein production by auto-induction in high density shaking cultures. (1046-5928 (Print)).

38. Dunne, O., et al., Matchout deuterium labelling of proteins for small-angle neutron scattering studies using prokaryotic and eukaryotic expression systems and high cell-density cultures. Eur Biophys J, 2017. 46(5): p. 425–432.

39. Haertlein, M., et al., Biomolecular Deuteration for Neutron Structural Biology and Dynamics. Methods Enzymol, 2016. 566: p. 113–57.

40. Tropea, J.E., S. Cherry, and D.S. Waugh, Expression and Purification of Soluble His6-Tagged TEV Protease, in High Throughput Protein Expression and Purification: Methods and Protocols, S.A. Doyle, Editor. 2009, Humana Press: Totowa, NJ. p. 297–307.

41. Muthana, M.M., et al., Improved one-pot multienzyme (OPME) systems for synthesizing UDP-uronic acids and glucuronides. Chem Commun (Camb), 2015. 51(22): p. 4595–8.

42. Nuell, M.J., et al., Prohibitin, an evolutionarily conserved intracellular protein that blocks DNA synthesis in normal fibroblasts and HeLa cells. Mol Cell Biol, 1991. 11(3): p. 1372–81.

43. Hu N-J., C.M., Daley M. & Hofmann A., Two new software applications for automated processing of circular dichroism and fluorescence data*. .* Appl Spectrosc 2005. 59: p. 68A.

44. SYSTAT, *SigmaPlot*. 2001, Systat Software, Inc.: San Jose, USA.

45. Micsonai, A., et al., BeStSel: a web server for accurate protein secondary structure prediction and fold recognition from the circular dichroism spectra. Nucleic Acids Res, 2018. 46(W1): p. W315–w322.

46. Lieutenant, K., P. Lindner, and R. Gahler, A new design for the standard pinhole small-angle neutron scattering instrument D11. Journal of Applied Crystallography, 2007. 40(6): p. 1056–1063.

47. Tully, M.D., et al., BioSAXS at European Synchrotron Radiation Facility - Extremely Brilliant Source: BM29 with an upgraded source, detector, robot, sample environment, data collection and analysis software. J Synchrotron Radiat, 2023. 30(Pt 1): p. 258–266.

48. Manalastas-Cantos, K.A.-O., et al., ATSAS 3.0: expanded functionality and new tools for small-angle scattering data analysis. (0021-8898 (Print)).

49. Svergun, D.I., Determination of the regularization parameter in indirect-transform methods using perceptual criteria. J. Appl. Crystallogr, 1992. 25: p. 495–503.

50. Pedersen, J., A flux- and background-optimized version of the NanoSTAR small-angle X-ray scattering camera for solution scattering. Journal of Applied Crystallography, 2004. 37(3): p. 369–380.

51. Glatter, O., A new method for the evaluation of small-angle scattering data. Journal of Applied Crystallography, 1977. 10(5): p. 415–421.

52. Pedersen, J.S., S. Hansen, and R. Bauer, The aggregation behavior of zinc-free insulin studied by small-angle neutron scattering. Eur Biophys J, 1994. 22(6): p. 379–89.

53. Bærentsen, R.L., et al., Structural basis for kinase inhibition in the tripartite E. coli HipBST toxin-antitoxin system. Elife, 2023. 12.

54. Dimasi, N., et al., *Expression,* crystallization and X-ray data collection from microcrystals of the extracellular domain of the human inhibitory receptor expressed on myeloid cells IREM-1. Acta Crystallographica Section F, 2007. 63(3): p. 204–208.

55. Zander, U., et al., Automated harvesting and processing of protein crystals through laser photoablation. Acta Crystallographica Section D, 2016. 72(4): p. 454–466.

56. Winter, G., et al., DIALS as a toolkit. Protein Science, 2022. 31(1): p. 232–250.

57. Winter, G., xia2: an expert system for macromolecular crystallography data reduction. Journal of Applied Crystallography, 2010. 43(1): p. 186–190.

58. Kabsch, W., XDS. Acta Crystallogr D Biol Crystallogr, 2010. 66(Pt 2): p. 125–32.

59. Evans, P.R., An introduction to data reduction: space-group determination, scaling and intensity statistics. Acta Crystallogr D Biol Crystallogr, 2011. 67(Pt 4): p. 282–92.

60. McCoy, A.J., et al., Phaser crystallographic software. J Appl Crystallogr, 2007. 40(Pt 4): p. 658–674.

61. Abramson, J., et al., Accurate structure prediction of biomolecular interactions with AlphaFold 3. Nature, 2024. 630(8016): p. 493-500.

62. Murshudov, G.N., A.A. Vagin, and E.J. Dodson, Refinement of macromolecular structures by the maximum-likelihood method. Acta Crystallogr D Biol Crystallogr, 1997. 53(Pt 3): p. 240–55.

63. Emsley, P., et al., Features and development of Coot. Acta Crystallogr D Biol Crystallogr, 2010. 66(Pt 4): p. 486–501.

64. Laskowski, R.A., et al., PROCHECK: a program to check the stereochemical quality of protein structures. Journal of Applied Crystallography, 1993. 26(2): p. 283–291.

65. Jones, D.T., Taylor, W. R., Thornton, J. M., A model recognition approach to the prediction of all-helical membrane protein structure and topology. Biochemistry, 1994. 33(10): p. 3038–3049.

66. Gao, K., R. Oerlemans, and M.A.-O. Groves, Theory and applications of differential scanning fluorimetry in early-stage drug discovery. (1867–2450 (Print)).

67. Micsonai, A., et al., BeStSel: a web server for accurate protein secondary structure prediction and fold recognition from the circular dichroism spectra. (1362–4962 (Electronic)).

68. Grudinin, S., M. Garkavenko, and A. Kazennov, Pepsi-SAXS: an adaptive method for rapid and accurate computation of small-angle X-ray scattering profiles. (2059–7983 (Electronic)).

69. Semenyuk, A.V. and D.I. Svergun, GNOM - a program package for small-angle scattering data processing. Journal of Applied Crystallography, 1991. 24(5): p. 537–540.

70. Honorato, R.V., et al., The HADDOCK2.4 web server for integrative modeling of biomolecular complexes. Nat Protoc, 2024. 19(11): p. 3219–3241.

71. Jumper, J., et al., Highly accurate protein structure prediction with AlphaFold.

72. Steglich, G., W. Neupert, and T. Langer, Prohibitins regulate membrane protein degradation by the m-AAA protease in mitochondria. Mol Cell Biol, 1999. 19(5): p. 3435–42.

73. Qiao, Z., et al., Cryo-EM structure of the entire FtsH-HflKC AAA protease complex. Cell Reports, 2022. 39(9): p. 110890.

74. Assar, Z., et al., Domain-Swapped Dimers of Intracellular Lipid-Binding Proteins: Evidence for Ordered Folding Intermediates. Structure, 2016. 24(9): p. 1590–1598.

75. Mayer-Harnisch, C.E., et al., N-terminal domain swapping: A new paradigm for spermidine/spermine N-acetyltransferase (SSAT) protein structures? Biochemical and Biophysical Research Communications, 2025. 748: p. 151302.

76. Duvezin-Caubet, S., et al., OPA1 processing reconstituted in yeast depends on the subunit composition of the m-AAA protease in mitochondria. Mol Biol Cell, 2007. 18(9): p. 3582–90.

77. Sacco, T., et al., Mouse brain expression patterns of Spg7, Afg3l1, and Afg3l2 transcripts, encoding for the mitochondrial m-AAA protease. BMC Neuroscience, 2010. 11(1): p. 55.

78. Göc, G., et al., Cryo-EM structure of the FtsH periplasmic domain reveals functional dynamics. bioRxiv, 2025: p. 2025.10.03.679995.doi: 10.1101/2025.10.03.679995.

79. Camargo, S., et al., Functional and structural characterization of an α-L-arabinofuranosidase from Thermothielavioides terrestris and its exquisite domain-swapped β-propeller fold crystal packing. Biochim Biophys Acta Proteins Proteom, 2020. 1868(12): p. 140533.

80. Asthana, P., et al., Structural insights into the substrate-binding proteins Mce1A and Mce4A from Mycobacterium tuberculosis. IUCrJ, 2021. 8(Pt 5): p. 757–774.

81. Mindrebo, J.T., et al., Cryo-EM Reveals Regulatory Mechanisms Governing Substrate Selection and Activation of Human LONP1. eLife, 2025. doi: 10.7554/eLife.109186.

